# The *C. elegans* DAF-19M module: a shift from general ciliogenesis to ciliary and behavioral specialization

**DOI:** 10.1101/2021.02.03.429678

**Authors:** Soungyub Ahn, Heeseung Yang, Sangwon Son, Dongjun Park, Hyunsoo Yim, Peter Swoboda, Junho Lee

**Author notes:** Co-corresponding authors, Junho Lee, Ph.D., Department of Biological Sciences, Seoul National University, Gwanak-ro 1, Gwanak-Gu, 08826, Seoul, Korea, Peter Swoboda, Ph.D., Department of Biosciences and Nutrition, Karolinska Institute, Campus Flemingsberg - NEO Building, Hälsovägen 9, SE-141 83 Huddinge, Sweden.

## Abstract

In animals, cilia are important for the interaction with environments and the proper function of tissues and organs. Understanding the distinctive identities of each type of ciliated cell is essential for therapeutic solutions for ciliopathies, complex disorders with impairments of various organs caused by defective cilia development and function. Here, we report a regulatory module consisting of a cascade of transcription factors and their target genes that confer the cell type-specific ciliary identities on the IL2 ciliated neurons in *C. elegans*. We found that DAF-19M, isoform of the sole *C. elegans* RFX transcription factor DAF-19, through X-box promoter motif variants, heads a regulatory module in IL2 neurons, comprising the core target genes *klp-6* (kinesin), *osm-9* (TRP channel), and *cwp-4* (novel); under the overall control of terminal selector proteins UNC-86 and CFI-1. Considering the conservation of this DAF-19M module in IL2 neurons for nictation, a dauer larva-specific behavior, and in male-specific neurons for mating behavior, we propose the existence of an evolutionarily adaptable, hard-wired genetic module for distinct behaviors that share the feature “recognizing the environment.”

## Introduction

The cilium is a microtubule-based structure that is anchored by the basal body (a modified centriole). The cilium protrudes from the cell surface like an antenna and thus, is a major cellular organelle for sensing the environment. Cilia can be categorized into two types: motile and non-motile cilia (also known as primary cilia). In humans, primary cilia sense and transmit signals to and from the immediate cell environment, while motile cilia provide rhythmic motion to move extracellular fluids or small particles across the cell surface (Anvarian *et al*, 2019; Mitchison & Valente, 2017; Nachury & Mick, 2019; Reiter & Leroux, 2017). Primary cilia have received widespread attention because they are present nearly ubiquitously on many different cell types in various tissues. Cilia regulate cellular processes in multiple organs, such as the brain, kidney, retina, liver, or the skeletal system (Mitchison & Valente, 2017). As such, sensing the environment by and information transfer through cilia are key cellular processes for the proper function of tissues and organs. Failure of cilia formation or proper cilia function can lead to a variety of ciliopathies, diseases in which the development and homeostasis of typically multiple organs are impaired. Ciliopathies that affect the nervous system result in distinct phenotypes and disorders, including brain malformations, cognitive impairment, and mental disorders of various strengths (Reiter & Leroux, 2017).

Because ciliopathies are often complex disorders and may present different phenotypes in different tissues, including in the nervous system, the study of individual or a small group of ciliated cells is essential for understanding exact molecular mechanisms and symptoms of a given ciliopathy. Utilizing an appropriate animal model is one way to overcome the inherent complexity of ciliopathies. The worm *Caenorhabditis elegans* is a useful model to study neuronal cilia thanks to the presence of ciliated sensory neurons and the overall simplicity of its nervous system (Inglis *et al*, 2007; White *et al*, 1986). Even though ciliated sensory neurons have the common identity to sense and transmit external signals through cilia, each one or even a small group of these neurons is also expected to have a unique identity to reflect its specific sensory modality and behavioral task, its interconnected position within the nervous system, and its spatiotemporal development and “final” position in the animal. The powerful genetics of *C. elegans* are ideally suited for the discovery of (upstream) regulators that determine the functional identity of an individual or a small group of ciliated sensory neurons (Etchberger *et al*, 2009; Etchberger *et al*, 2007; Inglis *et al*., 2007; Mukhopadhyay *et al*, 2007; Wang *et al*, 2010; Zhang *et al*, 2014).

To function as a sensory organelle and signal transducer, the fine-tuning of development and the maintenance of ciliary structure are essential. RFX transcription factors (TFs) are well known key regulators for the process of ciliogenesis (Choksi *et al*, 2014; Senti & Swoboda, 2008; Swoboda *et al*, 2000). The role of RFX TFs is to initiate and to maintain the regulation of genes for general ciliary structure and function; in a variety of cell types and tissues including in the nervous system; in a number of different organisms including in humans (Piasecki *et al*, 2010). Mammalian genomes encode RFX TF gene families (paralogs) and several of these paralogs play important roles during neuron development in several regions of the brain (Lemeille *et al*, 2020; Sugiaman-Trapman *et al*,2018). RFX TFs are defined by a certain type of DNA binding domain (DBD) and some have a dimerization domain (DIM) (Emery *et al*, 1996a; Gajiwala *et al*, 2000; Sugiaman-Trapman *et al*.,2018). RFX TFs recognize and bind to a *cis*-regulatory DNA sequence motif, the X-box motif, typically in promoters of direct target genes that are involved in the development and maintenance of cilia. X-box motif sequences are imperfect inverted repeats, consisting of two 6-nucleotides half sites separated by spacer nucleotides (GTNRCC N_1-3_ RGYAAC), to which RFX TFs can bind as homo-or hetero-dimers (Emery *et al*, 1996b; Gajiwala *et al*., 2000; Jolma *et al*, 2013).

This setup – RFX TF, X-box promoter motif, direct ciliary target gene – was first discovered in *C. elegans* (Swoboda *et al*., 2000). The *C. elegans* genome contains only one RFX TF gene, *daf-19*,which encodes several isoforms that govern different, yet in parts related biological functions. DAF-19A/B protein regulates synaptic homeostasis in non-ciliated neurons and DAF-19C regulates the developmental process of ciliogenesis in sensory neurons (De Stasio *et al*, 2018; Senti & Swoboda, 2008; Swoboda *et al*., 2000). Cilia and synapses share important conceptual and anatomical parallels in connection to their biological tasks as signal transducers (Shaham, 2010). DAF-19M regulates male mating behavior through a small group of specialized, male specific neurons that are ciliated and in which *daf-19m* is specifically expressed: CEM, RnB, and HOB (Wang *et al*., 2010). This particular group of ciliated neurons is thus a very good example for understanding how unique ciliary identities and functional specializations are attained, shaped and maintained.

Interestingly, *daf-19m* is also expressed in the six IL2 ciliated sensory neurons that are not sex-specific in the head of *C. elegans* (Wang *et al*., 2010). These neurons govern a very specific worm behavior, important for survival, nictation. Nictating worms wave their heads while standing on their tails (Cassada & Russell, 1975; Reed & Wallace, 1965). During the *C. elegans* life cycle, only dauer larvae, a juvenile diapause stage that is resistant to harsh environmental conditions, can nictate (Cassada & Russell, 1975). Nictation enables dauer larvae worms to latch on to carrier animals and get transported over long distances in search for more favorable environmental conditions (e.g. more food). In a previous study, we have determined that IL2 neurons, their intact, functional cilia and cholinergic neurotransmission are essential for nictation behavior (Lee *et al*, 2012). For all ciliated sensory neurons, including the IL2 neurons, general ciliary identity and functionality is regulated by DAF-19C, while the regulation of IL2 specific ciliary identity and specialization in connection to one dedicated behavior, nictation, has not been identified yet.

In this study, we have adopted a genetic screen to identify mutant animals with defective expression of an IL2 neuron specific identity gene, *klp-6*, encoding a ciliary kinesin (Peden & Barr, 2005). We found that *daf-19m* encodes a regulator of IL2 neuron specific ciliary identity, but not of non-ciliary identity features or characteristics. DAF-19M protein heads a functional module or regulatory subroutine comprising direct IL2 neuron ciliary target genes like *klp-6* (kinesin) or *osm-9* (TRPV channel) (Colbert *et al*, 1997; Peden & Barr, 2005; Wang *et al*., 2010). The proper function of this DAF-19M module is essential for the IL2 neuron controlled behavioral output, nictation. DAF-19M exerts its regulation by recognizing and binding to novel X-box motif variant sequences in the promoters of its direct target genes, while *daf-19m* itself is regulated by upstream terminal selectors, the TFs UNC-86 and CFI-1, which govern a number of aspects of terminal, functional differentiation of IL2 neurons (Zhang *et al*., 2014). Finally, we have used our in-depth knowledge of this regulatory cascade, the IL2 neuron DAF-19M module regulating direct targets through X-box motif variants, to search and find *C. elegans* genome-wide a large number of the novel, additional candidate genes that may contribute to generating the behavioral output of this module, nictation. The molecular identities of this novel set of genes will allow uncovering the mechanistic underpinnings of nictation.

## Results

### The gene *daf-19* encodes a regulator protein for IL2 neuron identity and functionality

The genes *klp-6* and *osm-3* encode kinesin motor proteins that are important for sensory cilia structure and function in IL2 and male specific neurons (Morsci & Barr, 2011; Peden & Barr, 2005). While – in both sexes – *osm-3* is expressed in many different types of ciliated sensory neurons including IL2 neurons, *klp-6* is specifically expressed only in IL2 neurons. In males, *klp-6* is also expressed in certain male specific ciliated sensory neurons. *klp-6* can thus be considered as an IL2 neuron identity gene (Wang *et al*., 2010; Zhang *et al*., 2014). Mutations in *klp-6* cause defects in the sensory behavior of nictation, governed by the IL2 neurons (Lee *et al*., 2012).

To find upstream regulators of IL2 neuron identity, we performed a forward genetic screen using a *klp-6* transcriptional GFP fusion (*klp-6p::gfp*) as a marker for IL2 neurons. We treated worms from the transgene insertion line *jlIs1900* with the mutagen <u>Et</u>hyl <u>m</u>ethane <u>s</u>ulfonate (EMS). The line *jlIs1900* carries both *klp-6p::gfp* and *aqp-6p::dsRed*, a marker for IL1 neurons. During development, IL1 and IL2 neurons differentiate from a common progenitor (Sulston *et al*, 1983). Therefore, to exclude mutations affecting the common progenitor of IL1 and IL2 neurons, we focused on isolating mutants with strongly decreased *klp-6p::gfp* expression in IL2 neurons that at the same time maintain intact *aqp-6p::dsRed* expression in IL1 neurons. We isolated EMS treated mutant worms using the COPAS BIOSORT (Fig 1A). From a screen of >15,000 F1 animals, we were able to isolate and maintain two mutations, both of which locate to the gene *daf-19* (Fig 1B): the *of3* allele locates to the dimerization domain (DIM), while the *of4* allele locates to the DNA binding domain (DBD) (De Stasio *et al*., 2018; Swoboda *et al*., 2000; Wang *et al*., 2010). Both mutations *of3* and *of4* cause strongly decreased *klp-6* expression (Fig 1C), fluorescent dye filling defects (Fig 1E) and a dauer constitutive phenotype (Daf-c). In genetic rescue experiments with a transgenic fosmid construct carrying wild-type *daf-19* (*jlEx1902*), all of these *daf-19* mutant phenotypes were restored, including prominent *klp-6* expression (Fig 1D and E). Additionally, we were able to recapitulate decreased *klp-6* expression in other *daf-19* mutant backgrounds (Fig EV1A). Importantly, IL2 neurons differentiate normally and their cell bodies and neurites are intact in a *daf-19* mutant background. Using a transcriptional GFP fusion to the *daf-19* independent IL2 gene *F28A12.3* as a marker *(F28A12.3p::gfp)* (Phirke *et al*, 2011), we could demonstrate intact expression in IL2 cell bodies and neurites (Fig EV1B), while cilia structure was still disrupted in a *daf-19(rh1024)* mutant background as these animals were still fluorescent dye filling defective.

**Figure 1.**
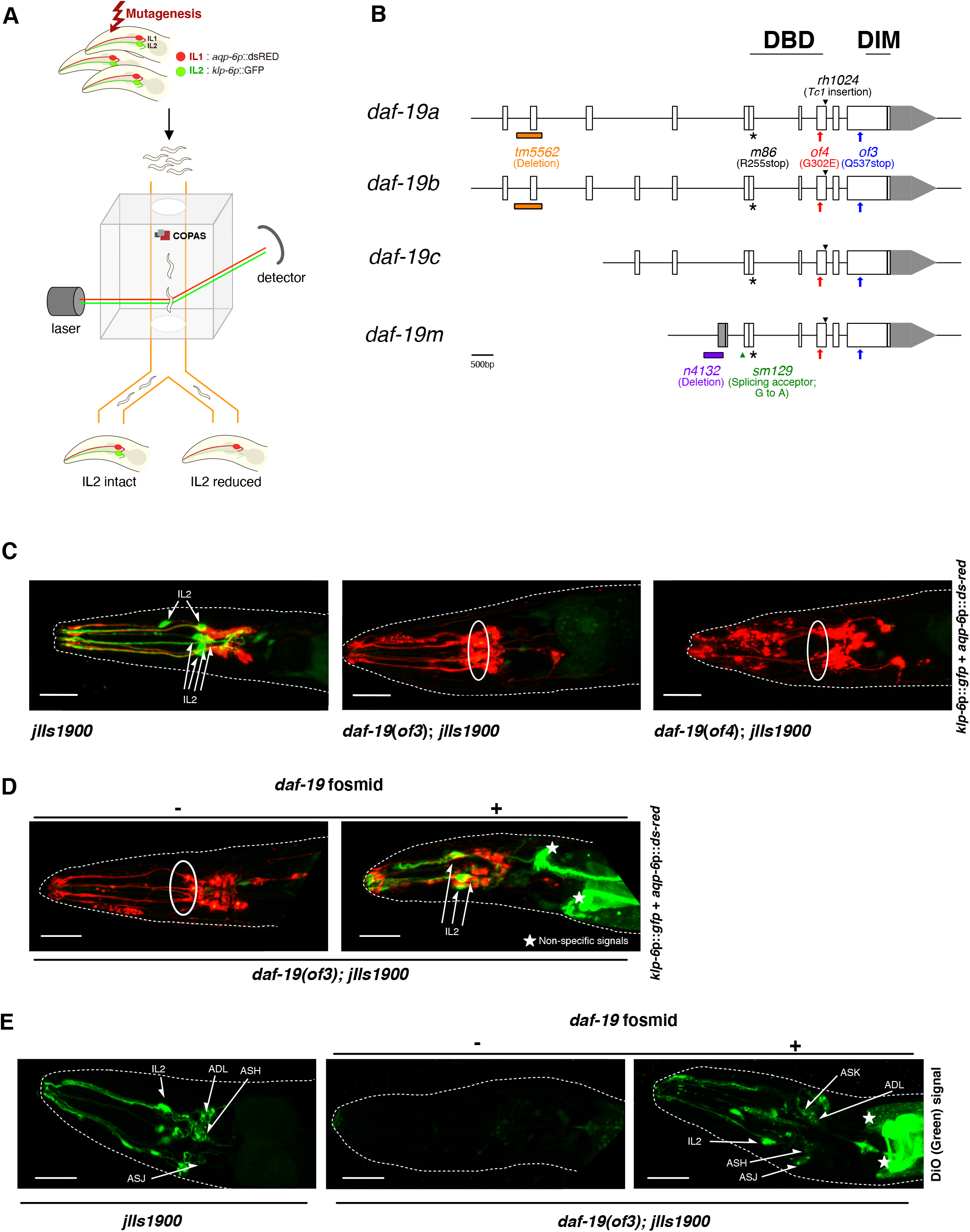
Novel mutations of the DAF-19/RFX transcription factor impact the expression of the IL2 neuron identity gene *klp-6*. **A** EMS mutagenesis scheme of *jlIs1900*, an integrated transgene carrying *klp-6p::gfp* and *aqp-6p::dsRed* as IL2 and IL1 neuronal markers, respectively. Mutants with decreased *klp-6* specific expression were isolated using the COPAS Biosort Flow Cytometer. **B** The *daf-19* specific mutant alleles *of3* and *of4* are located in the dimerization domain (DIM) and the DNA binding domain (DBD), respectively. The *of3* mutation affects amino acid 537 (Glutamine to Stop). The *of4* mutation affects amino acid 302 (Glycine to Glutamate). Other *daf-19* specific mutations relevant for this work are also indicated in the schematic. **C** Confocal images of *jlIs1900, daf-19(of3); jlIs1900*, and *daf-19(of4); jlIs1900*. In every image, the dotted line outlines the shape of the worm. IL2 neurons are indicated in the *jlIs1900* image (arrows). Solid-line ellipses indicate absent *klp-6p::gfp* fluorescent signals in IL2 neurons in both *daf-19(of3); jlIs1900* and *daf-19(of4); jlIs1900*. Scale bars are 20 μm. **D** Confocal images of *daf-19(of3); jlIs1900* with or without the presence of a transgene carrying functional *daf-19 (Ex[daf-19fosmid])*. In every image, the dotted line outlines the shape of the worm. IL2 neurons are indicated in the *daf-19(of3); jlIs1900; Ex[daf-19* fosmid] image (arrows). The solid-line ellipse indicates absent *klp-6p::gfp* fluorescent signals in IL2 neurons in *daf-19(of3); jlIs1900*. Non-specific signals are indicated in *daf-19 (of3)*; *jlIs1900; Ex*[*daf-19* fosmid] (Stars). Scale bars are 20 μm. **E** Confocal images of *jlIs1900* and *daf-19(of3); jlIs1900*, and *daf-19(of3)*; *jlIs1900*; *Ex*[*daf-19* fosmid] stained with the green-fluorescent dye DiO. In every image, the dotted line outlines the shape of the worm. Ciliated neurons that stain with DiO (IL2, ADL, ASH, ASJ, ASK) are indicated (arrows). Non-specific signals are indicated in *daf-19(of3); jlIs1900; Ex*[*daf-19* fosmid] (Stars). Scale bars are 20 μm.

In summary, we conclude that the new *daf-19* mutations *of3* and *of4* affect the expression of *klp-6* in IL2 neurons, IL2 sensory cilia development, and dauer formation, but not IL2 neuronal cell fate and its general differentiation into a neuron.

### Identification of a *cis*-regulatory element, an X-box motif variant, controlling *klp-6* expression

RFX transcription factors (TFs), including *C. elegans* DAF-19 protein, regulate gene expression by binding to X-box promoter motifs. However, a previous study reported that the *klp-6* promoter region lacks a canonical X-box motif (Fig EV2A) (Peden & Barr, 2005). Therefore, we systematically dissected the *klp-6* promoter region to find a *cis*-regulatory, presumed binding target for DAF-19 regulation. We created a series of *klp-6* promoter deletions and truncations that we fused to GFP as a marker for IL2 neurons in transgenesis experiments (Fig 2A). The overlapping transgene constructs *klp-6p (−614, −1)::gfp* and *klp-6p (−1048, −600)::gfp* showed intact expression in IL2 neurons indicating that the minimal region from −614 to −600 of the *klp-6* promoter is highly relevant for proper *klp-6* expression. Intriguingly, this region is similar to canonical X-box motifs, yet displays distinct deviations from canonical *C. elegans* X-box motif sequences (Fig 2B, EV2A and B, and Table EV2 and EV3; see also Materials and Methods). Therefore, we call this *klp-6* promoter *cis*-regulatory element an X-box motif variant.

**Figure 2.**
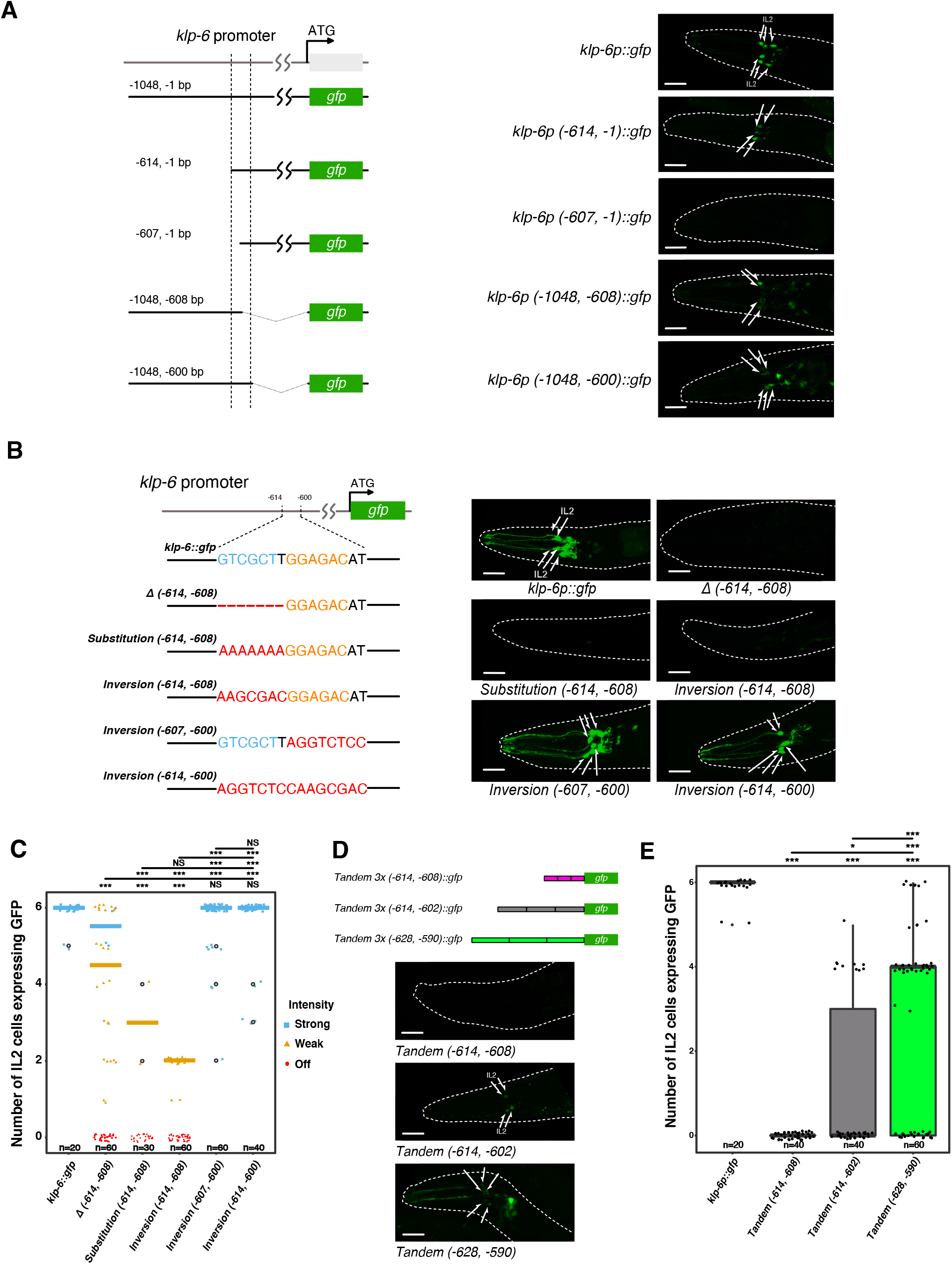
Identification of a *cis*-regulatory promoter element, an X-box motif variant, that controls *klp-6* expression in IL2 neurons. **A** *klp-6* promoter deletion analyses and confocal images of *Ex[klp-6p::gfp]* transgenic animals carrying the indicated *klp-6* promoter truncation construct. *klp-6* promoter deletion analyses identified the relevant promoter region that controls *klp-6* expression in IL2 neurons. Truncated *klp-6* promoters were fused to the GFP gene using the fusion PCR method. In every image, the dotted line outlines the shape of the worm. IL2 neurons are indicated (arrows). Scale bars are 20 μm. **B** Illustration of mutagenesis of the *klp-6* promoter and confocal images of *Ex*[*klp-6p::gfp*] transgenic animals carrying the indicated *klp-6* promoter construct mutation. Mutagenesis of the *klp-6* promoter determined the necessity of the relevant promoter region, an X-box motif variant, for *klp-6* expression in IL2 neurons: deletion *(Δ −614, −608:* from −614 to −608 upstream of the *klp-6* start codon ATG); substitution (sequences from −614 to −608 were substituted with AAAAAAA); and three different inversions (sequences from −614 to −608 or from −607 to −600 or from −614 to −600 were substituted with the corresponding sequences from the complementary strand). These *klp-6* promoter mutations were carried out using a commercial plasmid mutagenesis kit in a *klp-6p::gfp* plasmid background, itself based on the vector pPD114.108. In every image, the dotted line outlines the shape of the worm. IL2 neurons are indicated (arrows). Scale bars are 20 μm. **C** Quantification of GFP expression in transgenic animals carrying *Ex[klp-6p::gfp]* with the indicated *klp-6* promoter construct mutation. 20 worms per independent transgenic line were examined except for the *substitution (−614, −608)* line, where 15 worms were examined. We distinguished between strong, weak and absent (off) GFP expression and determined in how many of the six IL2 neurons GFP expression was detected. Statistical significance was determined with one-way ANOVA and Tukey’s multiple comparison post-test. *** *P*≤0.001, significantly different from the control. NS, not significant. Overall *p* value for ANOVA is less than 0.001 (*P*<0.001). **D** Illustration of tandem repeats of *klp-6* X-box motif variant minimal promoter regions and confocal images of transgenic animals carrying the indicated tandem repeat of the *klp-6* X-box motif variant minimal promoter regions fused to GFP: 5’ half site motif, full motif, extended motif. Tandem repeats of three different *klp-6* X-box motif variants fused to GFP; magenta – 5’ half of the X-box motif variant (−614, −608); grey – full X-box motif variant (−614, −602); light green – extended X-box motif variant (−628, −590) including adjacent promoter sequences that contain one additional 5’ and 3’ X-box like half site, respectively (see also Fig EV2A). The different tandem repeats were created *de novo* in an empty GFP vector (pPD95.77) background using a commercial plasmid mutagenesis kit. In every image, the dotted line outlines the shape of the worm. IL2 neurons are indicated (arrows). Scale bars are 20 μm. **E** Quantification of GFP expression in transgenic animals carrying the indicated tandem repeat of the *klp-6* X-box motif variant minimal promoter regions fused to GFP: 5’ half site motif, full motif, extended motif. 20 worms per independent transgenic line were examined. We determined in how many of the six IL2 neurons GFP expression was detected. Statistical significance was determined with one-way ANOVA and Tukey’s multiple comparison post-test. * *P*≤0.5, *** *P*≤0.001, significantly different from the control. NS, not significant. Overall *p* value for ANOVA is less than 0.001 (*P*<0.001).

We mutagenized this X-box motif variant to investigate in transgenesis experiments its necessity for *klp-6* expression (Fig 2B and C). When we interfered with the 5’ half site of the X-box motif variant, *klp-6* expression decreased drastically. For example, only 1 of 3 transgenic lines of the *Δ (−614, −608)* construct showed weak *klp-6* expression in some IL2 neurons, while in 2 of 3 lines *klp-6* expression was absent. In contrast, interfering with the 3’ half site did not significantly affect *klp-6* expression in IL2 neurons. Only some transgenic animals showed expression in a reduced number of IL2 neurons, while most of the transgenic animals showed entirely intact expression. Importantly, the overall intensity of *klp-6* expression remained uniformly strong. We conclude that the 5’ half site of the X-box motif variant is more critical for *klp-6* expression than its 3’ half counterpart. An inversion of the full *klp-6* X-box motif variant did not affect *klp-6* expression in IL2 neurons, as also the *klp-6* X-box motif variant, like canonical X-box motifs, is an imperfect inverted repeat sequence.

We then cloned tandem repeats of the *klp-6* X-box motif variant into the pPD95.77 GFP vector to investigate in transgenesis experiments whether this motif is also sufficient for *klp-6* expression (Fig 2D and E). Tandem repeats of the 5’ half site *(Tandem 3x (−614, −608)::gfp)* did not elicit GFP expression in IL2 neurons. In contrast, tandem repeats of the full X-box motif variant *(Tandem 3x (−614, −602)::gfp)* or of an extended X-box motif variant *(Tandem 3x (−628, −590)::gfp*) were able to elicit (often weak) GFP expression in IL2 neurons. We noted that in some cases the extended X-box motif variant was able to elicit entirely intact GFP expression in all six IL2 neurons (Fig 2E).

Like other genes that are (specifically) expressed in IL2 neurons in both sexes, *klp-6* is also expressed in the CEM, HOB and RnB male specific neurons (Peden & Barr, 2005; Wang *et al*.,2010). And like is the case for IL2 neurons, the minimal region from −614 to −600 of the *klp-6* promoter was necessary for male specific neuronal expression of *klp-6* (Fig EV2C). Likewise, a transgene construct carrying the extended X-box motif variant *(Tandem 3x (−628, −590)::gfp)* was able to elicit (often weak) GFP expression in the CEM, HOB and RnB neurons (Fig EV2D).

Together, these results demonstrate and support the conclusion that the *klp-6* X-box motif variant is both necessary and sufficient for regulating gene expression not only in IL2 but also in male specific neurons. Accordingly, and underscoring its importance for regulating *klp-6* expression, the *klp-6* X-box motif variant is well conserved in other *Caenorhabditis* species like in *C. briggsae* and in *C. brenneri* (Fig EV2B).

### The protein isoform DAF-19M regulates genes in IL2 neurons through the X-box motif variant

The expression of *klp-6* is restricted to IL2 and male specific neurons (Peden & Barr, 2005). To investigate whether DAF-19 regulates *klp-6* expression in IL2 neurons by itself or together with co-factor(s), we carried out a yeast-1-hybrid (Y1H) screening experiment. Using the extended *klp-6* promoter X-box motif variant sequence as bait, we isolated 119 independent yeast colonies, each representing an independent binding event. 114 of 119 yeast colonies carried cDNA constructs that resulted in DAF-19 protein binding to the bait sequence (Table EV1). We confirmed the molecular identity of the underlying cDNA constructs from all 119 colonies in all cases by PCR and in a few cases by sequencing. We note that all presently known isoforms of DAF-19 (A/B, C, M) share exactly the same DNA binding domain (DBD) (De Stasio *et al*., 2018; Senti & Swoboda, 2008; Swoboda *et al*., 2000; Wang *et al*., 2010). This heterologous Y1H experiment does therefore not allow distinguishing which isoform binds to the *klp-6* promoter X-box motif variant. Our Y1H results indicate, however, that in all likelihood DAF-19 regulates *klp-6* expression without a co-factor.

*klp-6p::gfp* transgene expression is strongly decreased or absent in *daf-19* mutant backgrounds that affect all the isoforms, or affect the *daf-19c* or the *daf-19m* isoforms, but is entirely intact in a *daf-19a/b* specific mutant background (Fig EV1A) (Wang *et al*., 2010). Similar observations were made for other genes specifically expressed in IL2 neurons (De Stasio *et al*., 2018; Wang *et al*., 2010). To investigate which DAF-19 isoform regulates gene expression in IL2 neurons through the *klp-6* promoter X-box motif variant, we carried out transgenic rescue experiments determining GFP expression of the extended X-box motif variant construct *(Tandem 3x (−628, −590)::gfp)* (Fig 3A and B, EV4A). A transgene construct containing *daf-19c* was not able to restore GFP expression, while a construct containing *daf-19m* did restore GFP expression in IL2 neurons, albeit only partially. Providing both *daf-19m* and *daf-19c* restored GFP expression even further, nearly to wild-type levels, both with regard to IL2 cell numbers and GFP intensity (Fig 3A and B). We conclude that primarily the isoform DAF-19M is necessary for IL2 neurons gene expression through the X-box motif variant, while DAF-19C provides supporting function.

**Figure 3.**
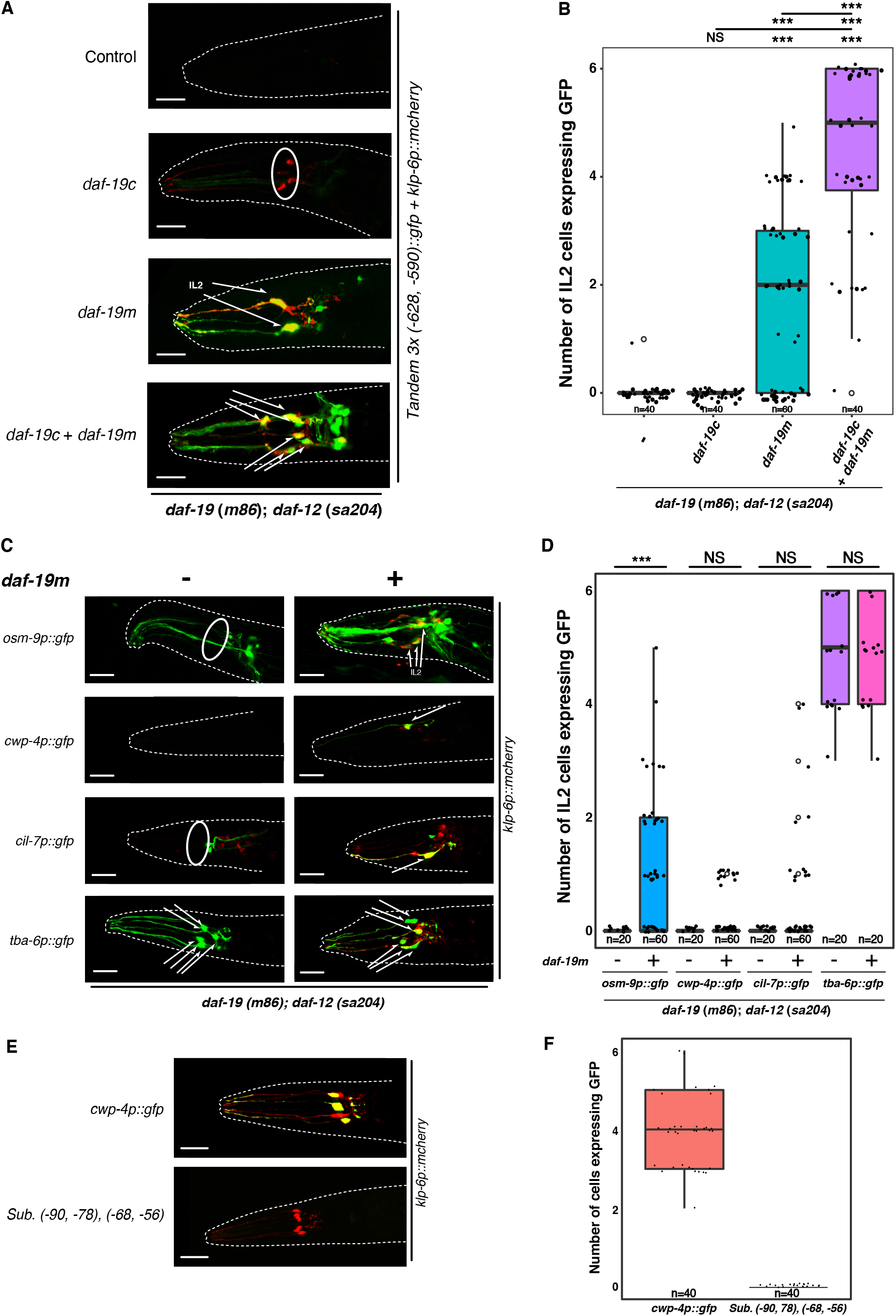
The gene isoform *daf-19m*, originally discovered for male mating, regulates genes expressed in IL2 neurons through an X-box promoter motif variant. **A** Confocal images of *daf-19* isoform specific rescue in a *daf-19* null mutant background [JT6924: *daf-19 (m86); daf-12(sa204)]*. We revealed genetic rescue by using the tandem repeat construct with the *klp-6* extended X-box motif variant minimal promoter region fused to GFP. Constructs for *daf-19* isoform specific rescue are as follows: *pGG14* and *pJL1920* for *daf-19c; pJL1921* for *daf-19m; pGG14* and *pJL1921* for *daf-19c* and *daf-19m* (see also Fig EV4A). In every image, the dotted line outlines the shape of the worm. IL2 neurons are indicated (arrows), whereby *klp-6p::mCherry* was used as an IL2 neuron specific marker. In one image, the solid-line ellipse indicates *klp-6p::mCherry* red-fluorescent signals in unidentified, but not IL2, neurons. Scale bars are 20 μm. **B** Quantification of GFP expression in transgenic animals carrying the indicated *daf-19* isoform specific rescue construct. 20 worms per independent transgenic line were examined. We determined in how many of the six IL2 neurons GFP expression was detected. Statistical significance was determined with one-way ANOVA and Tukey’s multiple comparison post-test. *** *P*≤0.001, significantly different from the control. Overall *p* value for ANOVA is less than 0.001 (*P*<0.001). **C** Confocal images of transgenic animals carrying promoter-to-GFP fusions of four genes prominently expressed and functioning in IL2 neurons: *osm-9, cwp-4, cil-7, tba-6*. We determined gene expression in a *daf-19* null mutant background [JT6924: *daf-19(m86); daf-12 (sa204)*], without (-) or with (+) the addition of a *daf-19m* specific rescue construct. In every image, the dotted line outlines the shape of the worm. IL2 neurons are indicated (arrows), whereby *klp-6p::mCherry* was used as an IL2 neuron specific marker. Solid-line ellipses indicate absent *klp-6p* specific fluorescent signals in IL2 neurons. Scale bars are 20 μm. **D** Quantification of GFP expression in transgenic animals carrying promoter-to-GFP fusions for four IL2 neuron genes *(osm-9, cwp-4, cil-7, tba-6)* without (-) or with (+) the addition of a *daf-19m* specific rescue construct. 20 worms per independent transgenic line were examined. We determined in how many of the six IL2 neurons GFP expression was detected. Statistical significance was determined with one-way ANOVA and Tukey’s multiple comparison post-test. *** *P*≤0.001, significantly different from the control. NS, not significant. Overall *p* value for ANOVA is less than 0.001 (*P*<0.001). **E** Confocal images of *Ex[cwp-4p::gfp]* transgenic animals carrying either the wild-type *cwp-4* promoter or the indicated *cwp-4* promoter mutation construct. The *cwp-4* promoter mutation consisted of substituting both X-box variant sequences (−90 to −78 and −68 to −56) with GGATCC C GGATCC. These substitutions were carried out using a commercial plasmid mutagenesis kit in a fusion PCR generated *cwp-4p::gfp* background. In every image, the dotted line outlines the shape of the worm. IL2 neurons are indicated (arrows), whereby *klp-6p::mCherry* was used as an IL2 neuron specific marker. Scale bars are 20 μm. **F** Quantification of GFP expression in transgenic animals carrying either the wild-type *Ex*[*cwp-4p::gfp*] promoter construct or the indicated promoter construct mutation. 20 worms per independent transgenic line were examined. We determined in how many of the six IL2 neurons GFP expression was detected.

Next, we investigated whether other genes specifically expressed in IL2 neurons, and highly relevant for IL2 ciliary functionality, were also regulated by DAF-19M (Fig 3C and D). In previous studies, *osm-9* (encoding a ciliary TRPV channel), *cwp-4* (a novel gene encoding a protein with a PTS/mucin domain), *cil-7* (encoding a myristolated protein localizing to cilia), and *tba-6* (encoding an alpha-tubulin), have been reported to be expressed in IL2 and male specific neurons (Colbert *et al*., 1997; Hurd *et al*, 2010; Maguire *et al*, 2015; Portman & Emmons, 2004). The expression of *osm-9, cwp-4*, and *cil-7* was restored in IL2 neurons in transgenic *daf-19m* rescue experiments, while *tba-6* expression was *daf-19m* independent. *daf-19m* rescue of *osm-9* expression was strong, while rescue of *cwp-4* and *cil-7* expression, respectively, was weaker and not statistically significant: it allowed for only partial expression in some of the six IL2 neurons (Fig 3D).

To investigate whether the DAF-19M mediated regulation of this (mostly ciliary) gene expression occurs through an X-box motif variant as in *klp-6*, we searched for similar sequence motifs in the genes *osm-9, cwp-4*, and *cil-7*. Within and directly up-and downstream (5’ and 3’) of the gene *osm-9*,we were so far unable to find a sequence hit that resembles the *klp-6* X-box motif variant (Colbert *et al*., 1997). The promoter region of *cil-7* harbors such a sequence hit similar to *klp-6*, which, however, is not conserved in other *Caenorhabditis* species (from Wormbase; www.wormbase.org). Interestingly, a highly similar doublet of the *klp-6* X-box promoter motif variant exists in the *cwp-4* promoter region (Fig 3E): located at −90 to −78 and at −68 to −55 upstream of the start codon ATG. These *C. elegans cwp-4* X-box variants are well conserved in other *Caenorhabditis* species, like in the orthologous *cwp-4* promoter regions of *C. briggsae*, *C. remanei*, *C. brenneri*, and *C. japonica* (Fig EV3A). We mutated these X-box motif variant sequences in the *cwp-4* promoter and investigated in transgenesis experiments their necessity for *cwp-4* expression. We found that *cwp-4* expression was entirely absent upon motif mutation, both in IL2 neurons (Fig 3E and F) as well as in the CEM, HOB and RnB; male specific neurons (Fig EV3B). We have thereby confirmed the concept of DAF-19M regulating gene expression through an X-box motif variant for at least two genes, *klp-6* and *cwp-4*.

### DAF-19M heads a regulatory subroutine in IL2 neurons

Terminal selectors, which initiate the terminal differentiation of neurons, confer and maintain neuronal identities (Hobert, 2011, 2016). The genes *unc-86* and *cfi-1* encode such terminal selectors that confer and maintain specific identities in IL2 neurons, like being cholinergic or being ciliated sensory neurons (Zhang *et al*., 2014). Since DAF-19M regulates a number of genes important for proper IL2 function, we wondered whether the gene *daf-19m* itself is a member of the IL2 neuron terminal selector group headed by UNC-86 and CFI-1, and thus, whether expression of *daf-19m* is dependent on these terminal selectors. The expression of *klp-6* was reduced significantly in IL2 terminal selector mutant backgrounds (Zhang *et al*., 2014). And similar to *klp-6*, the expression of *daf-19m* was absent in *unc-86*(*n846*), and strongly reduced in *cfi-1*(*ky651*) mutant backgrounds (Fig 4A and B).

**Figure 4.**
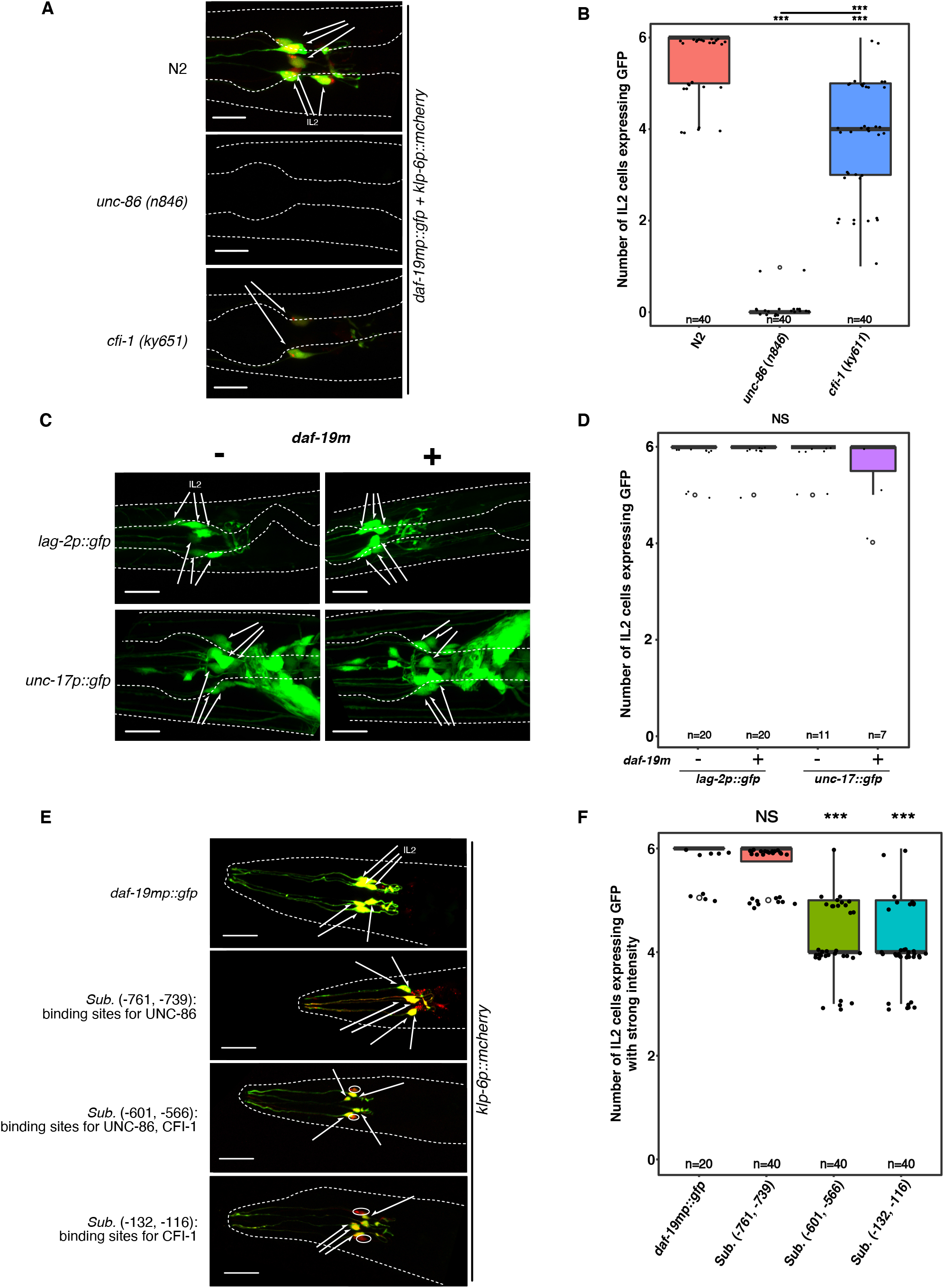
The gene isoform *daf-19m* is regulated by IL2 neuron terminal selectors and comprises a regulatory subroutine with its downstream genes. **A** Confocal images of *Ex*[*daf-19mp::gfp*] transgenic animals in a wild-type N2 background or in IL2 neuron terminal selector mutant, *unc-86(n846)* and *cfi-1(ky651)*, backgrounds. In every image, the outer dotted line outlines the shape of the worm and the inner dotted line outlines parts of the pharynx. IL2 neurons are indicated (arrows), whereby *klp-6p::mCherry* was used as an IL2 neuron specific marker. Scale bars are 10 μm. **B** Quantification of GFP expression in transgenic animals carrying *Ex[daf-19mp::gfp]* in the indicated backgrounds. 20 worms per independent transgenic line were examined. We determined in how many of the six IL2 neurons GFP expression was detected. Statistical significance was determined with one-way ANOVA and Tukey’s multiple comparison post-test. *** *P*≤0.001, significantly different from the control. NS, not significant. Overall *p* value for ANOVA is less than 0.001 (*P*<0.001). **C** Confocal images of transgenic animals carrying promoter-to-GFP fusions of two IL2 neuron identity genes: *lag-2p::gfp (qIs56)* and *unc-17p::gfp (vsIs48)* in a wild-type N2 and in a *daf-19m* specific mutant background [*daf-19(n4132)*]. In every image, the outer dotted line outlines the shape of the worm and the inner dotted line outlines parts of the pharynx. IL2 neurons are indicated (arrows). Scale bars are 10 μm. **D** Quantification of GFP expression in transgenic animals carrying promoter-to-GFP fusions of IL2 neuron identity genes (*lag-2, unc-17*) in the indicated backgrounds: wild type *versus daf-19m* specific mutant [*daf-19(n4132)*]. 20 worms per independent transgenic line were examined for *lag-2p::gfp*,while 7 and 11 worms, respectively, were examined for *unc-17p::gfp*. We determined in how many of the six IL2 neurons GFP expression was detected. Statistical significance was determined with one-way ANOVA and Tukey’s multiple comparison post-test. NS, not significant. Overall *p* value for ANOVA is 0.2169. **E** Confocal images of Ex[*daf-19mp::gfp*] transgenic animals carrying either a wild-type *daf-19m* promoter construct or constructs where different candidate binding sites for IL2 neuron terminal selector proteins, UNC-86 and CFI-1, have been mutated by substitution with poly-A sequences as indicated (see also Fig EV4B). In every image, the dotted line outlines the shape of the worm. IL2 neurons are indicated (arrows), whereby *klp-6p::mCherry* was used as an IL2 neuron specific marker. Solid-line ellipses indicate absent *daf-19mp::gfp* fluorescent signals in IL2 neurons marked by *klp-6p::mCherry*. Scale bars are 20 μm. **F** Quantification of GFP expression with strong intensity in transgenic animals carrying either a wild-type Ex[*daf-19mp::gfp*] promoter construct or constructs with mutations of the indicated candidate binding sites for IL2 neuron terminal selector proteins, UNC-86 and CFI-1. 20 worms per independent transgenic line were examined. We determined in how many of the six IL2 neurons GFP expression of mutated *daf-19m* promoter was detected with strong intensity, compared to wild-type Ex[*daf-19mp::gfp*]. Statistical significance was determined with one-way ANOVA and Tukey’s multiple comparison post-test. *** *P*≤0.001, significantly different from the control. NS, not significant. Overall *p* value for ANOVA is less than 0.001 (*P*<0.001).

We then examined two IL2 identity genes that are part of the same UNC-86 and CFI-1 terminal selector group, but whose gene products are not connected to IL2 ciliary functionality: *lag-2* (encoding a notch receptor ligand that is expressed in IL2 neurons in dauer larvae) and *unc-17* (encoding a protein involved in IL2 cholinergic synaptic transmission) (Ouellet *et al*, 2008; Rand, 1989; Zhang *et al*., 2014). The expression of *lag-2* and *unc-17* was unchanged between wild type and *daf-19(n4132)*, the *daf-19m* specific mutant background (Fig 4C and D). These results suggest that within the UNC-86 and CFI-1 terminal selector group, DAF-19M heads a regulatory subroutine consisting of IL2 identity genes with relevant ciliary functionality (Fig 3C and D), while non-ciliary IL2 identity genes are *daf-19m* independent (Fig 4C and D).

A regulatory subroutine can be regarded as a defined module, which as an entity forms part of a terminal selector group. Such a subroutine module consists of a TF (which itself may directly be regulated by the terminal selectors heading the group) and its direct downstream or effector genes, which then confer specific neuronal identities to terminally differentiating neurons (Altun-Gultekin *et al*, 2001; Etchberger *et al*., 2007; Gordon & Hobert, 2015; Hobert, 2008, 2016). We examined whether *daf-19m* is directly regulated by the terminal selectors and TFs UNC-86 and CFI-1. We first determined candidate binding sites for both UNC-86 and CFI-1 in the *daf-19m* promoter region (Fig EV4B). We then generated substitution mutations in defined candidate UNC-86 and CFI-1 binding sites in a *daf-19mp::gfp* expression construct and examined the resulting transgenic GFP expression patterns in IL2 neurons. Only mutations in certain binding sites for both UNC-86 and CFI-1 (−601 to −566; −132 to −116) reduced the GFP expression in IL2 neurons: typically four instead of six IL2 neurons still expressed GFP with strong intensity (Fig 4E and F), thereby partially phenocopying *daf-19m* expression in a *cfi-1* mutant background (Fig 4A and B) (Zhang *et al*., 2014). These results suggest that both multiple UNC-86 and CFI-1 binding sites (−601 to −566) and multiple CFI-1 binding sites (−132 to −116) are essential for *daf-19m* expression.

DAF-19M is thus the TF heading a regulatory subroutine for IL2 neuron ciliary functionality and is directly regulated by the terminal selectors and TFs UNC-86 and CFI-1. We note that the DAF-19M regulatory subroutine appears to maintain activity even after (embryonic) IL2 neuron development and differentiation is complete, as *daf-19m* and its downstream genes continue to be prominently expressed in IL2 neurons post-development and differentiation during the L1 and dauer larval stages (Fig EV1C and EV3C).

### The DAF-19M regulatory subroutine controls nictation behavior

In a previous study, we reported that cilia structure and function, as well as synaptic transmission in IL2 neurons, are essential for nictation behavior (Lee *et al*., 2012). In particular, *klp-6* and *osm-9* mutants that have defective IL2 cilia structure, show reduced nictation ratios. DAF-19M regulates *klp-6* and *osm-9* in IL2 neurons (Fig 3C and D) (Wang *et al*., 2010), and thus, we hypothesized that mutations in *daf-19m* would also impact nictation behavior.

In a series of experiments, we determined nictation ratios for various *daf-19* mutants and for mutants in IL2 genes with relevant ciliary functionality (Fig 5). Unfortunately, it proved technically impossible to determine nictation ratios for *daf-19(m86)*, the null mutant affecting all *daf-19* isoforms, as *daf-19(m86)* dauers move very little. *daf-19(tm5562)*, a *daf-19a/b* isoform specific mutant, did not show a significant nictation defect (Fig 5A). Of note, in the *daf-19(tm5562)* mutant background the expression of *klp-6* is fully intact (Fig EV1A). On the other hand, *daf-19(n4132) and daf-19(sm129)*,two different *daf-19m* isoform specific mutants, showed clear nictation defects (Fig 5A and B). Next, in genetic rescue experiments, we attempted to restore the nictation defects of the *daf-19m* isoform specific mutant, *daf-19(n4132)*. Providing *daf-19m* as a transgene was not sufficient to rescue the *daf-19(n4132)* nictation defects, while providing the direct *daf-19m* downstream genes *osm-9, klp-6* and *cwp-4* as transgenes did enable the rescue of nictation defects (Fig 5C). We then examined mutants of other direct *daf-19m* downstream genes (Fig 5D and E): *cwp-4(tm727)* and *osm-9(yz6)* mutants clearly showed significantly reduced nictation ratios, while *cil-7(tm5848)* mutants did not. *cil-7* mutants only showed nictation defects in a *cil-7; klp-6* double mutant background as both *cil-7 (tm5848); klp-6(ys71)* and *cil-7(tm5848); klp-6(ys72)* showed significantly reduced nictation ratios.

**Figure 5.**
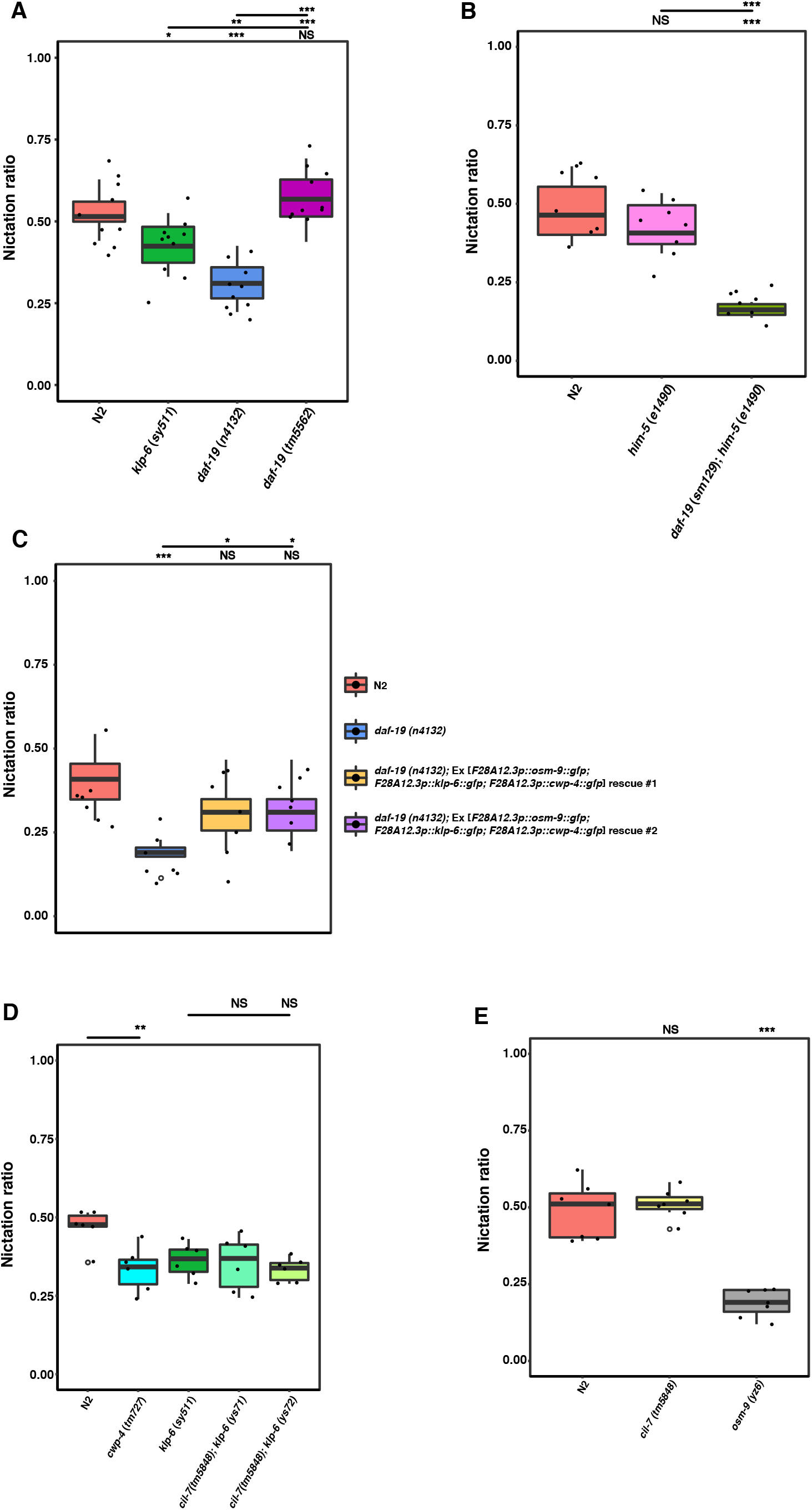
The *daf-19m* regulatory subroutine regulates *C. elegans* nictation behavior. **A** Nictation ratios of wild-type N2, and the mutants *klp-6(sy511), daf-19(n4132)* – a *daf-19m* isoform specific mutant, and *daf-19(tm5562)* – a *daf-19a/b* isoform specific mutant. Statistical significance was determined with one-way ANOVA and Tukey’s multiple comparison post-test. * *P*≤0.5, ** *P*≤0.1, *** *P*≤0.001, significantly different from the control. NS, not significant. Overall *p* value for ANOVA is less than 0.001 (*P*<0.001). **B** Nictation ratios of wild-type N2, and the mutants *him-5(e1490)*, and *daf-19 (sm129)* – a mutant where a splice acceptor site for the *daf-19m* isoform is disrupted – in a *him-5(e1490)* background. Statistical significance was determined with one-way ANOVA and Tukey’s multiple comparison post-test. *** *P*≤0.001, significantly different from the control. Overall*p* value for ANOVA is less than 0.001 (*P*<0.001). **C** Nictation ratios of wild-type N2, and the mutant *daf-19(n4132)* – a *daf-19m* isoform specific mutant, and of transgenic lines expressing rescuing genomic DNA constructs of *osm-9, klp-6*, and *cwp-4*, downstream target genes of *daf-19m* functioning in IL2 neurons. Statistical significance was determined with one-way ANOVA and Tukey’s multiple comparison post-test. * *P*≤0.5, *** *P*≤0.001, significantly different from the control. NS, not significant. Overall *p* value for ANOVA is less than 0.001. **D** Nictation ratios of wild-type N2, and mutants of *daf-19m* downstream target genes that are expressed and functional in IL2 neurons: *cwp-4(tm727), klp-6(sy511)*, and *cil-7 (tm5848); klp-6(ys71)*and *cil-7(tm5848); klp-6(ys72)*. Statistical significance was determined with one-way ANOVA and Tukey’s multiple comparison post-test. ** *P*≤0.1, significantly different from the control. NS, not significant. Overall *p* value for ANOVA is 0.0058. **E** Nictation ratios of wild-type N2, and mutants of genes that are expressed and functional in IL2 neurons with previously demonstrated roles in IL2 sensory cilia exposed to the environment at the tip of the head of the worm: *cil-7(tm5848)* and *osm-9(yz6)*. Statistical significance was determined with one-way ANOVA and Tukey’s multiple comparison post-test. *** *P*≤0.001, significantly different from the control. NS, not significant. Overall *p* value for ANOVA is less than 0.001 (*P*<0.001).

We conclude that *klp-6, osm-9* and *cwp-4*, direct *daf-19m* downstream genes in IL2 neurons, encode key proteins highly relevant for nictation behavior, strongly suggesting that the DAF-19M regulatory subroutine is crucial for enabling nictation behavior through IL2 neuron function.

### Predicting additional DAF-19M downstream genes with an X-box promoter motif variant

We hypothesized that additional and novel genes, candidates for also being involved in nictation governed by IL2 neurons, might also be regulated by DAF-19M through an X-box motif variant. To search for such additional and novel members of the DAF-19M subroutine throughout the *C. elegans* genome, we used the *klp-6* and *cwp-4* X-box motif variants and very similar sequences as search tools (see Materials and Methods for details). We built position weight matrices (PWMs) of both X-box motif variants and of *C. elegans* canonical X-box motifs for reference, including in both cases a number of experimentally proven X-box motifs (Fig 2B, EV2A and B, EV3A, Table EV2 and EV3). We used both PWMs to carry out searches throughout the *C. elegans* genome. In these genome-wide searches, we allowed the 1-3 spacer nucleotide(s) to be N, NN or NNN, as in a variety of species, including in *C. elegans*, the (vast) majority of X-box motifs contains a double nucleotide spacer, while single and triple nucleotide spacer sequences have also been found. As expected in all our genome-wide searches (given the complexities of the respective query sequence PWMs and the overall similarities versus differences between canonical X-box motifs and X-box motif variants), we found large numbers of X-box candidate hits (>75.000). In particular in searches using NN double nucleotide spacers, the search output contained a number of hits of already experimentally proven X-box motifs of ciliary genes, strongly validating our overall search strategy (Table EV4, EV5, and EV6). To focus on candidate X-box hits as part of a DAF-19M regulatory subroutine in IL2 neurons, we sorted and filtered for potential promoter motifs, by firstly requiring the candidate motif hit to locate within 1 kb of the nearest start codon ATG of a given gene and secondly requiring these candidate genes to have a suspected or experimentally demonstrated expression pattern that included IL2 neurons (as extracted from Wormbase, version WS235; www.wormbase.org). The search results then yielded 62 genes for the X-box motif variant PWM with a single spacer nucleotide N as query, 97 genes for the PWM with double spacer nucleotides NN, and 66 genes for the PWM with triple spacer nucleotides NNN (Table EV4, EV5, and EV6). We anticipate that these gene lists contain significant numbers of novel candidate members of the DAF-19M regulatory subroutine in IL2 neurons. Ongoing and future work will determine DAF-19M dependent gene expression in IL2 neurons, mutant analyses involvement in IL2 governed nictation, and molecular characterization of encoded protein function will provide mechanistic insight into how nictation behavior is generated and then executed by the worm.

## Discussion

RFX transcription factors (TFs), by binding to X-box motifs in promoters of general ciliary genes, regulate the developmental process of ciliogenesis (Choksi *et al*., 2014). First demonstrated for DAF-19, the *C. elegans* RFX TF (Swoboda *et al*., 2000), this finding was subsequently replicated in many different organisms, including mammals (Choksi *et al*., 2014; Piasecki *et al*., 2010). The gene *daf-19* encodes several different protein isoforms that regulate different biological functions in addition to ciliogenesis (De Stasio *et al*., 2018; Senti & Swoboda, 2008; Wang *et al*., 2010; Wells *et al*, 2015; Xie *et al*, 2013). In this study, we have focused on the role of the protein isoform DAF-19M, which heads a functional module (or regulatory subroutine) controlling organismal behaviors, like male mating (Wang *et al*., 2010) and nictation (this work). Interestingly, the regulation of this DAF-19M module goes through an X-box motif variant, which mediates the expression in IL2 neurons of crucially important genes for nictation, like *klp-6* and *cwp-4*. X-box motifs are imperfect inverted 6-nucleotides repeat sequences separated by 1-3 spacer nucleotides (Blacque *et al*, 2005; Chen *et al*, 2006; Efimenko *et al*, 2005; Emery *et al*., 1996b). Very often the X-box spacer nucleotides are AT, as is the case for most *C. elegans* ciliary genes regulated by the ciliogenic DAF-19C protein isoform (Blacque *et al*., 2005; Burghoorn *et al*, 2012; Efimenko *et al*., 2005). Here, we have uncovered a shorter X-box motif variant with only a 1-nucleotide spacer T. Also, compared to canonical *C. elegans* X-boxes, this variant appears to be less stringently conserved at crucial motif positions (Gajiwala *et al*., 2000) as well as with regard to positioning within promoter regions upstream of gene starting codons (ATG) (Burghoorn *et al*., 2012). These aspects may have precluded the discovery of such an X-box motif variant in previous, quite stringently constructed X-box motif search efforts (Blacque *et al*., 2005; Burghoorn *et al*., 2012; Chen *et al*., 2006; Efimenko *et al*., 2005). Using expression assays employing *klp-6* and *cwp-4*, two genes crucial for ciliary functionality of IL2 neurons, we have shown that the X-box motif variant is both necessary and sufficient for expression in IL2 neurons, and in male specific neurons. Thereby, the X-box motif variant proves essential for nictation behavior and possibly also for male mating. It thus provides an important entry point into uncovering the molecular mechanisms that govern behaviors like nictation and male mating through functional modules like the one headed by DAF-19M (Fig 6).

**Figure 6.**
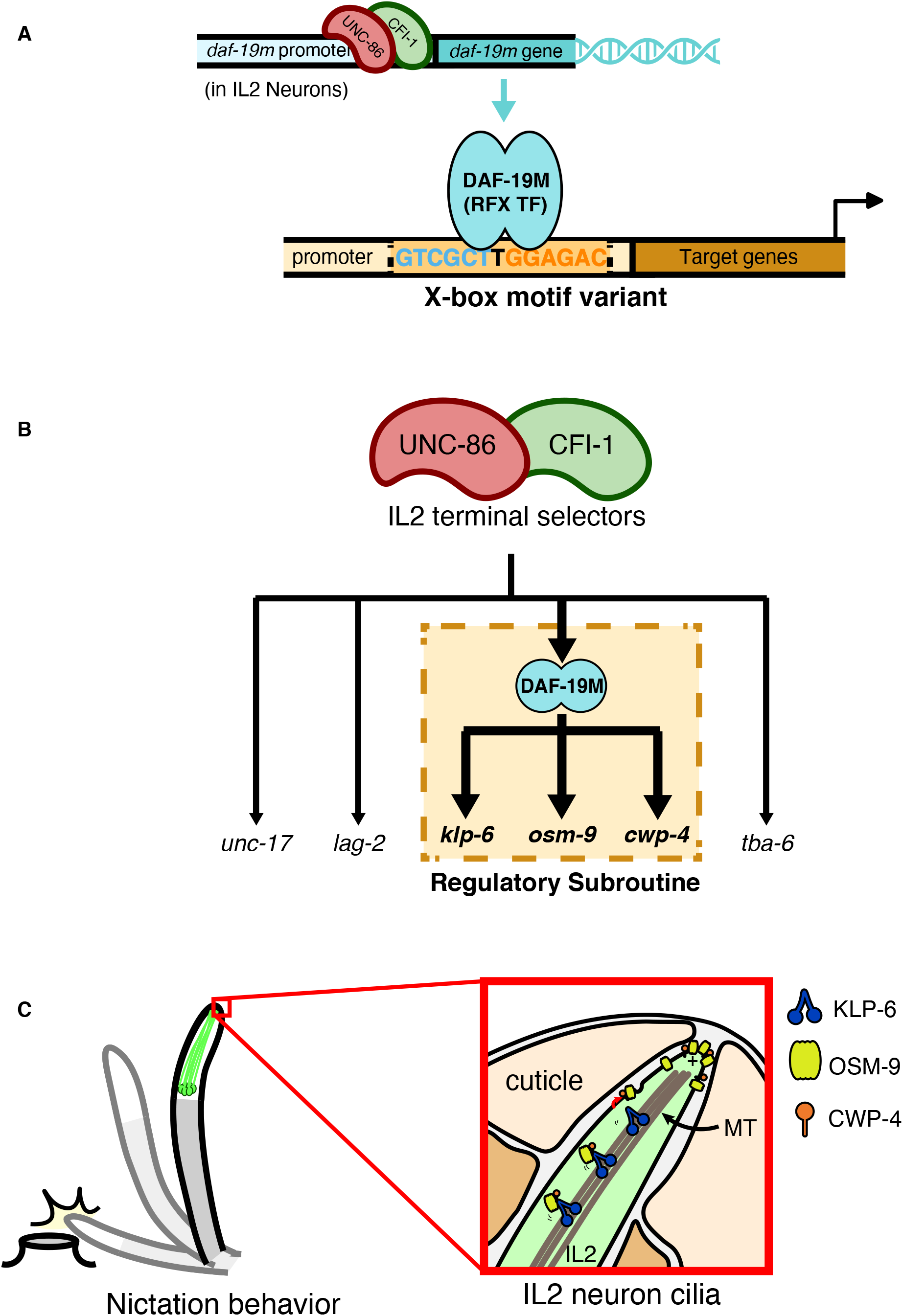
A model describing the importance of the *daf-19m* regulatory subroutine for IL2 neuron function. **A** DAF-19M protein, regulated by IL2 neuron terminal selector proteins UNC-86 and CFI-1, controls the expression of its target genes through an X-box promoter motif variant. **B** We have experimentally demonstrated that DAF-19M protein heads a regulatory subroutine that comprises at least three (but very likely more) genes prominently expressed and functioning in IL2 neurons and their sensory cilia: *klp-6*, *osm-9*, and *cwp-4*. **C** The *daf-19m* regulatory subroutine controls nictation behavior through the function of IL2 neurons. The KLP-6 kinesin transports proteins relevant and necessary for proper nictation to the ciliary tip of IL2 neurons. IL2 sensory cilia are exposed to the environment at the tip of the worm’s head.

Our work also features a general aspect that may be highly relevant for other experimental systems. We have demonstrated that small shifts in TF binding motif sequence conservation or lack thereof, from canonical X-box motif to X-box motif variant, can have functional consequences, from governing general ciliary functionality to cell-specific ciliary specializations and behavioral output. These shifts may thus provide molecular mechanisms for how to adopt new biological functions step-by-step.

How does DAF-19M through binding to the X-box motif variant regulate the expression of its target genes in IL2 neurons? Our Y1H results do not distinguish which of the DAF-19 protein isoforms binds to the X-box motif variant sequence. DAF-19 protein isoforms differ only by their N-terminal amino acid sequences, while central and C-terminal sequences, including crucial functional domains like DBD and DIM are identical between all the isoforms (Senti & Swoboda, 2008; Wang *et al*., 2010). In the IL2 ciliated neurons, both gene isoforms *daf-19c* and *daf-19m* are expressed, while *daf-19a/b* is expressed in the nervous system specifically in non-ciliated neurons (De Stasio *et al*., 2018; Senti & Swoboda, 2008; Wang *et al*., 2010). DAF-19C and DAF-19M differ by only a few N-terminal amino acids, while the C-terminal 611 amino acids are identical. DAF-19M has 11 N-terminal amino acids not shared with DAF-19C, while DAF-19C has 50 or 27 N-terminal amino acids not shared with DAF-19M (depending on starting in exon 4 or 5, respectively) (from Wormbase; www.wormbase.org). In our genetic rescue experiments, we have determined that in IL2 neurons DAF-19M can regulate the expression of its ciliary target genes *klp-6* and *cwp-4*, while DAF-19C by itself cannot. This DAF-19M mediated rescue is incomplete though, but can be elevated to nearly wild-type levels by supplying also DAF-19C. This indicates that in IL2 neurons DAF-19M can bind to X-box motif variant sequences as homo-dimer, albeit inefficiently, while DAF-19C supported hetero-dimers with DAF-19M elevate this binding efficiency to (functionally) nearly wild-type levels. How then is binding accomplished and distinguished, respectively, between canonical X-boxes in promoters of general ciliary genes (DAF-19C targets) and X-box motif variants in promoters of genes for functional ciliary specializations like *klp-6* and *cwp-4* (primarily targets of DAF-19M supported by DAF-19C), given that both types of target genes, and both *daf-19c* and *daf-19m*, are all expressed in IL2 ciliated neurons? One possibility is that the (slight) differences in N-terminal amino acids between DAF-19C and DAF-19M might affect 3-D protein (TF) structure and thereby impact binding affinities to canonical X-boxes versus X-box motif variants. Alternatively, and possibly more likely, IL2 neuron specific protein co-factors (activators and/or inhibitors) might impact the ability of binding to canonical X-boxes versus X-box motif variants, respectively. Such a co-factor scenario has been proposed for the impact DAF-19 has on the serotonin neurotransmitter biosynthesis pathway in the ADF ciliated neurons, even though in that case no X-boxes or variants have hitherto been found in the promoters of serotonin pathway genes (Xie *et al*., 2013). Co-factors might also be involved in modulating the activities of RFX TF paralogs in mammals, where it was found in the mouse that the RFX1-3 paralogs can regulate the same or different sets of (ciliary) target genes depending on cellular context, but without dedicated specificity between the respective RFX TF paralog (1, 2 or 3) and the sequence composition of a given X-box motif (Lemeille *et al*., 2020). In the future, it will be interesting to examine the interactions between DAF-19M and DAF-19C (and X-box motif variants) using microscopy-based techniques such as BiFC, BRET or *in vitro* affinity assays (Bhuckory *et al*, 2019; Lai & Chiang, 2013), so as to gain mechanistic insight into how the IL2 neuron specific DAF-19M functional module initiates and operates.

The TFs UNC-86 and CFI-1 have previously been described as terminal selectors for IL2 neurons as they confer, regulate and maintain specific (anatomical and molecular) identities that are characteristic for terminally differentiated, functional IL2 neurons (Zhang *et al*., 2014). We have shown that DAF-19M is a constituent of the UNC-86/CFI-1 terminal selector group. Both TFs directly regulate DAF-19M through defined binding sites in the *daf-19m* promoter. In turn, DAF-19M through the X-box motif variant then directly regulates its IL2 neuron specific target genes and thereby heads a regulatory subroutine within the UNC-86/CFI-1 terminal selector group. Given the molecular identities of DAF-19M and its direct target genes this regulatory subroutine can be defined as a functional module concerning IL2 neuron ciliary functionality (Fig 6A and B).

What is the biological role of the DAF-19M regulatory subroutine (or functional module) in IL2 neurons considering the molecular identities and functions of the DAF-19M direct target genes? *klp-6* encodes a specialized ciliary kinesin motor protein, which is expressed in IL2 and some male specific neurons. KLP-6 has been shown to transport TRP channels in male specific neurons (Morsci & Barr, 2011; Peden & Barr, 2005), while its cargo in IL2 neurons has not been reported yet. Interestingly, *osm-9* encodes a neuronal TRPV channel that regulates chemotaxis, osmotic avoidance, and touch response. *osm-9* is expressed in ciliated sensory neurons, including the IL2 neurons (Colbert *et al*.,1997; Wang *et al*., 2010). *cwp-4* encodes a novel ciliary protein that is expressed in IL2 and some male specific neurons. *cwp-4* encodes an N-terminal signal peptide for secretion and a C-terminal membrane anchoring domain (Miller & Portman, 2010; Portman & Emmons, 2004). We speculate that CWP-4 protein is held in place in the ciliary membrane and functions outside the cell (cilium), where it may be available for protein interaction or as a co-factor for cell (ciliary) surface channel or receptor proteins. We propose therefore that in IL2 ciliated neurons the DAF-19M functional module serves to transport, set and keep in place a molecular (protein) machinery for chemo-and/or mechano-sensation: e.g. receptors, channels, extra-and intra-cellular co-factors and signal transducers; possibly even at the ciliary tip that is directly exposed to the environment (Fig 6C). Such a setup would ensure that nictation, an essential behavior for worm survival, can properly be carried out (Lee *et al*, 2017; Lee *et al*., 2012). IL2 neurons would thereby be able to sense and transduce the relevant external stimuli for nictation: (i) sense the surface environment for when and where to initiate nictation; and (ii) sense and interact with potential animal carriers to initiate the hitchhiking behavior for the worm to be passively transported over long distances in search for more suitable environmental conditions (e.g. more or better food) that improve chances for survival. It will in the future be interesting to determine the exact subcellular or ciliary localizations of the proteins involved in nictation, by using in transgenic rescue experiments targeted translational GFP fusion proteins that examine relevant protein domains. Also, measuring neuronal activity of IL2 neurons upon physical stimulation using calcium imaging will enable to test our hypothesis concerning DAF-19M and its direct target genes.

We have uncovered that the DAF-19M functional module also operates in *C. elegans* male specific ciliated neurons. In male specific neurons, DAF-19M regulates the ciliary kinesin gene *klp-6* (and also *cwp-4*), and thereby the TRP channels LOV-1 and PKD-2, which are KLP-6 cargoes (Morsci & Barr, 2011; Peden & Barr, 2005; Wang *et al*., 2010). Both TRP channels localize to cilia in neurons of the male tail. Both TRP channels and KLP-6 are involved in sensing hermaphrodites during male mating behavior, which requires crucial chemo-and mechano-sensory steps (Barr & Garcia, 2006; Barr *et al*, 2018), striking parallels to the above described nictation behavior. We speculate that nictation and male mating, two distinct yet mechanistically similar behaviors in connection to the sensation and recognition of the environment (nictation – the surface and a carrier animal; mating – the mating partner), are regulated by the same program: the DAF-19M functional module. This module is genetically hard-wired given that both behaviors are essential for worm survival, at the individual level (nictation) and at the population level (male mating). The DAF-19M module might provide a template for how to organize at the molecular level these essential biological functions. We provide experimental evidence for the first three direct targets of DAF-19M –*klp-6, cwp-4* and *osm-9* – and present their importance for nictation (for their potential relevance and impact on male mating behavior see: Miller & Portman, 2010; Peden & Barr, 2005; Zhang *et al*, 2018). We predict the molecular machineries governing sensory aspects of both behaviors to be complex and depending on (changing) environmental conditions to be heavily tunable. Thus, we expect a substantially larger set of direct DAF-19M target genes. To this end, we have generated genome-wide *bona fide* DAF-19M candidate target gene lists based on the presence of high confidence promoter X-box motif variant sequence hits and (highly likely) expression in IL2 neurons. Cross-comparing our genome-wide work with other efforts that have yielded genes functioning in IL2 neurons (Wang *et al*, 2015) will greatly facilitate (i) extracting new members of the DAF-19M functional module and then (ii) uncovering and analyzing in detail its mechanistic impact on nictation and male mating.

## Materials and Methods

### Maintenance of *C. elegans* worms and worm strains used

All worm strains were maintained at 20°C and handled as previously described (Brenner, 1974), except for strains carrying *daf-19(of3), daf-19(of4), daf-19(m86)*, or *daf-19(rh1024)* mutant alleles, as these cause high frequency of dauer larva formation at 20°C (Daf-c phenotype). These *daf-19* mutant strains were grown at 15°C instead (Swoboda *et al*., 2000). Some *C. elegans* strains were obtained from the *Caenorhabditis* Genetics Center (CGC, University of Minnesota, St. Paul, MN, USA; https://cgc.umn.edu) or from the National BioResource Project, Japan (https://shigen.nig.ac.jp/c.elegans). See Table EV7 for a complete list of mutant and transgenic strains that were used in this study.

### Generation of transgene constructs and of transgenic animals

PCR-based GFP-fusion constructions were carried out for *klp-6* promoter deletion analysis (Hobert, 2002). Both *klp-6p::gfp* and *klp-6p::mCherry* fusion constructs were created using classical restriction enzyme-based subcloning methods. Other constructs were created using a Gibson assembly cloning kit (E5510; New England Biolabs). All DNA fragments were inserted into the GFP vector backbone pPD95.77, except for *klp-6p::gfp* (inserted into pPD114.108) and *klp-6p::mCherry* (inserted into pPD117.01; modified to *mCherry* red fluorescence). Plasmid constructs were modified by site-directed mutagenesis (E0554; New England Biolabs) to create the desired mutations in *klp-6* and *cwp-4*, X-box promoter motif variant sequences (Fig 2C: deletions, substitutions, inversions; 2F: insertions; 3F: substitutions) and in the *daf-19m* promoter in candidate binding sites for IL2 neuron terminal selector proteins UNC-86 and CFI-1 (Fig 4E: substitutions). All the cloning and assembly details of all transgene constructs are available on request.

To generate transgenic animals, we microinjected DNA and plasmid constructs into the gonad of young adult hermaphrodites as previously described (Mello *et al*, 1991). To isolate transgenic progeny of microinjected hermaphrodites, we used the following co-injection markers for transgenesis: *rol-6 (su1006sd)* (roller phenotype), *unc-122p::dsRed* (red fluorescent coelomocytes), *myo-2p::mCherry* (red fluorescent pharynx), and *act-5p::gfp* (green fluorescent intestine). Plasmid DNAs used for microinjection were extracted and purified with a QIAGEN plasmid midi kit (Cat. No. 12145) or an Axygen midi prep kit (Cat. No. AP-MD-P-25) or a Macherey-Nagel NucleoBond Xtra Midi kit (Cat. No. 740410.100).

### Transgene insertion into the genome by gamma-ray irradiation

To facilitate our forward genetic screening approach, synchronized L4 larvae of the transgenic strain *jlEx1900* [*klp-6p::gfp; aqp-6p::dsRed; rol-6(su1006sd)*] were gamma-ray irradiated at 4000 rad. Irradiated worms were moved individually to new plates to determine whether the *jlEx1900* transgene was successfully integrated into the genome; by measuring the proportion of the roller phenotype in progeny worms. When the transgenesis marker (roller phenotype) had reached full penetrance (100%) on a given plate of progeny worms, expression of the relevant plasmid constructs, *klp-6p::gfp* in IL2 neurons and *aqp-6p::dsRed* in IL1 neurons, was confirmed by fluorescence microscopy. Only integrated lines with fully penetrant expression of both plasmid constructs and the transgenesis marker were used for follow-up outcrossing and mutagenesis work.

### Ethyl methane sulfonate (EMS) mutagenesis and isolation of mutant worms

To find regulators of *klp-6* expression in IL2 neurons, the strain LJ800: *jlIs1900* [*klp-6p::gfp; aqp-6p::dsRed; rol-6(su1006sd)*] was mutagenized with 50mM EMS and >15,000 worms of the F1 progeny generation were examined for GFP and DsRed expression changes. Mutations were identified by reduced *klp-6p::gfp* expression in IL2 neurons, both with regard to the number of IL2 neurons expressing GFP and overall GFP fluorescence intensity. Two independent mutant lines were isolated using the COPAS BIOSORT large particle flow cytometer (Union Biometrica, MA, USA) for high throughput sorting; measuring fluorescence intensity, time-of-flight (for animal length), and extinction (optical density). Through canonical SNP mapping and whole genome sequencing, we found that both mutations located in the gene *daf-19*. We named these mutant alleles *of3* and *of4*,respectively.

## Fluorescence microscopy and sample preparation for imaging

Confocal microscopy (ZEISS LSM700; Carl Zeiss) and fluorescence microscopy (Axioplan 2; Carl Zeiss) were used to observe transgene expression in IL2 neurons and – as needed – in other cell types, and for the acquisition of fluorescence images. For microscopy and imaging, transgenic animals were paralyzed with 3mM levamisole and mounted on 3% agar pads. All transgenic animals were observed and imaged during the first day of adulthood, except when other developmental and life stages were used for experimentation (Fig EV1A and 1C; strains LJ805 and LJ806).

## Dauer formation assays and fluorescent dye staining to ascertain mutations in the gene *daf-19*

Dauer formation assays and DiO (green fluorescent dye) staining were performed as previously described (Perkins *et al*, 1986; Starich *et al*, 1995). The green fluorescent dye DiO stains a small number of ciliated sensory neurons in the head and tail of *C. elegans*, all of which are directly exposed to the environment, including in the head the IL2 neurons (Schroeder *et al*, 2013; Ward *et al*, 1975; White *et al*., 1986).

## Induction of dauer formation

Ten L4 larvae or young adults within the first day of adulthood were moved to synthetic pheromone plates, including a thin layered lawn of *E. coli* OP50 bacteria, at 25°C for dauer induction (Lee *et al*, 2015; Lee *et al*., 2017; Lee *et al*., 2012). Synthetic pheromone plates contain agar (10g/L), agarose (7g/L), NaCl (2g/L), KH2PO4 (3g/L), K2HPO4 (0.5g/L), cholesterol (8mg/L) and the pheromones ascaroside 1, 2, 3 (2mg/L each) (Butcher *et al*, 2007; Jeong *et al*, 2005; Lee *et al*.,2017). Synthetic pheromones were provided by the Young-Ki Paik laboratory at Yonsei University, Seoul, Korea. After 4 to 5 days, dauer larvae are easily recognizable by their radially constricted bodies and their dark intestines. The induction of dauer formation was determined in populations consisting of at least 100 worms.

## Nictation assays

We first created a micro-dirt chip by pouring a 3.5% agar solution onto a PDMS mold (Lee *et al*.,2015). The solidified chip was then detached from the PDMS mold and dried for 90 min at 37°C. For nictation assays, more than 30 dauer larvae were collected by a glass capillary tube using M9 isotonic buffer and mounted onto a freshly prepared micro-dirt chip. After 10-30 min, when dauers started to move, nictation was quantified as the fraction of nictating worms among moving dauers (Lee *et al*., 2012). Quiescent dauers were excluded from measurements. Nictation assays were carried out at 25°C with a humidity of 30%. Assays were repeated at least six times for quantification and statistics.

## Analysis of gene expression in IL2 neurons

*C. elegans* has a total of six IL2 neurons in the head. We counted the number of IL2 neurons that express a given gene promoter-to-fluorescent marker fusion (GFP or mCherry) and – when needed – also determined the intensity of GFP or mCherry expression in IL2 neurons. For additional distinction, GFP or mCherry expression intensity was divided into three categories: strong, weak, and off (absent). At least 20 worms per transgenic line were examined except for one *klp-6p::gfp substitution (−614, −608)* line (Fig 2E) and the transgenic lines for examining *unc-17p::gfp* expression (Fig 4D). Typically, we examined GFP or mCherry expression in transgenic animals of at least two independent transgenic lines with same genetic background. For some experiments, we used *klp-6p::mCherry* as marker for IL2 neurons to confirm the correct expression in IL2 neurons of *daf-19m* and other IL2-expressed genes fused to GFP.

## Yeast-one-hybrid (Y1H) screening assays

The regulatory element of the *klp-6* promoter that we have identified (−628, −590; gtccgtttcc tttcgtcgct tggagaccta catggcaac) was cloned as target sequence or bait into the pADE2i vector, constructed by Panbionet (Pohang, Korea), for Y1H screening in order to identify candidate proteins that bind to this element. This vector, designed for the Matchmaker Gold Yeast One Hybrid library (Clontech, CA, USA), contains the yeast iso-1-cytochrome C minimal promoter and the ADE2 gene. As prey *C. elegans* cDNAs were inserted into the pPC86 vector, containing the GAL4 activation domain (AD). We used these GAL4 AD fusion libraries to screen for cDNAs encoding proteins that interact with the target or bait sequence. Positive interactions (positive years clones) showed ADE2 expression. We used PCR and sequencing to examine positive clones (Table EV1).

## Statistical analyses

For all experimental quantifications, the statistical significance was determined with one-way ANOVA and Tukey’s multiple comparison post-test, except for when a given class of results was zero and therefore statistics could not be applied (Fig 3G).

## Bioinformatics: genome-wide sequence motif searches

The X-box DNA sequence motif is an imperfect, inverted repeat, consisting of two 6-nucleotide half-sites (5’ and 3’) separated by 1-3 spacer nucleotide(s). Its canonical sequence composition is bound by RFX transcription factors (TFs) and has been determined both *in vitro* and *in vivo* (Emery *et al*., 1996b; Gajiwala *et al*., 2000; Jolma *et al*., 2013; Swoboda *et al*., 2000). The sole *C. elegans* RFX TF gene, *daf-19*, encodes different isoforms that all share the same DNA binding domain (DBD) (Fig 1B) (De Stasio *et al*.,2018; Senti & Swoboda, 2008; Swoboda *et al*., 2000; Wang *et al*., 2010). Through work on DAF-19 ciliary target genes, the sequence composition of the canonical X-box motif is very well known in worms (Table EV2 and EV3) (Blacque *et al*., 2005; Burghoorn *et al*., 2012; Chen *et al*., 2006; Efimenko *et al*., 2005).

The RFX TF isoform DAF-19M heads a regulatory subroutine in IL2 neurons (Fig 6), which employs an X-box motif variant that is slightly different from a canonical X-box motif (Fig 2 and Fig 3; Fig EV2; Table EV2 and EV3). To search for potentially additional members of this DAF-19M subroutine throughout the *C. elegans* genome, we used the founding member of this subroutine, the *klp-6* X-box motif variant and very similar sequences as a search tool.

First, we used the MEME software suite (version 5.2.0; http://meme-suite.org/tools/meme) to build position weight matrices (PWMs) of both X-box variants and of canonical X-boxes as reference. The *C. elegans* canonical X-box motif, with very rare exceptions (e.g. *nph-1;* Burghoorn *et al*., 2012), consists of two 6-nucleotides half sites (5’ and 3’) separated by a double nucleotide (AT) spacer, whereby positional nucleotide conservation reflects experimentally determined binding characteristics between the RFX TF (DAF-19) DBD and the X-box DNA sequence motif (Emery *et al*., 1996b; Gajiwala *et al*., 2000; Jolma *et al*., 2013; Swoboda *et al*., 2000). For example: (i) positions 1 and 3 of the 5’ X-box half site are most often G and T, and only in a minority of cases A and C, respectively; (ii) the double nucleotide spacer sequence is almost exclusively AT; (iii) positions 4 and 6 of the 3’ X-box half site are most often A and C, and only in a minority of cases G and T, respectively.

We built a PWM of the canonical X-box motif based on 40 X-box sequences derived from a number of *C. elegans* genes with experimentally proven X-boxes and from their direct orthologs in closely related *Caenorhabditis* species (Table EV2) (Blacque *et al*., 2005; Burghoorn *et al*., 2012; Chen *et al*., 2006; Efimenko *et al*., 2005). Of these 40 X-box sequences, 18 fit an “ideal” sequence conservation pattern (5’ half site position 1 = G and position 3 = T; double nucleotide spacer = AT; 3’ half site position 4 = A and position 6 = C), 21 have one deviation from an “ideal” sequence conservation pattern, while 1 has two deviations. We then built a PWM of X-box motif variants based on 37 X-box sequences derived from a mixture of *C. elegans* genes with experimentally proven X-boxes and of candidate X-box sequence hits, and from their direct orthologs in closely related *Caenorhabditis*species, respectively (Table EV3) (Blacque *et al*., 2005; Burghoorn *et al*., 2012; Chen *et al*., 2006; Efimenko *et al*., 2005; including this work). Of these 37 X-box sequences, 10 contain only a single nucleotide spacer like is the case for the *klp-6* X-box promoter motif variant. Of the remaining 27 X-box sequences with a double nucleotide spacer, only 4 fit an “ideal” sequence conservation pattern (5’ half site position 1 = G and position 3 = T; double nucleotide spacer = AT; 3’ half site position 4 = A and position 6 = C), while 17 have one deviation from an “ideal” sequence conservation pattern, and 6 have two deviations. It is apparent that both PWMs (canonical versus variant) share large overlaps yet also present slight differences (see also the sequence logos in Table EV2 and EV3), acknowledging that the different isoforms of the sole *C. elegans* RFX TF, DAF-19, share the exact same DBD.

Secondly, we used both PWMs to carry out genome-wide searches through the *C. elegans* genome. All candidate X-box sequence motif hits were extracted using the FIMO tool (Find Individual Motif Occurrences; version 5.3.0; http://meme-suite.org/tools/fimo), whereby the FIMO search parameter p-value was required to be smaller than 1E-04 (standard cut-off setting). We allowed the 1-3 spacer nucleotide(s) to be N, NN or NNN. This to reflect the fact that the *klp-6* X-box motif variant contains only a single nucleotide spacer (T) and that in a variety of species the majority of X-box motifs contain a double nucleotide spacer, while single and triple nucleotide spacer sequences have also been found (Burghoorn *et al*., 2012; Emery *et al*., 1996b; Laurençon *et al*, 2007; Piasecki *et al*., 2010; Sugiaman-Trapman *et al*., 2018).

As expected in all our genome-wide searches (given the lengths and complexities of query sequence PWMs; canonical versus variant; allowing for 1-3 spacer nucleotides), we found large numbers of X-box candidate hits (>75,000), whereby in searches using NN double nucleotide spacers the search output contained well-represented hits of already known and experimentally proven X-box motifs of ciliary genes (Table EV5).

Thirdly, to focus on candidate X-box hits as part of a DAF-19M regulatory subroutine in IL2 neurons, we employed the following sorting and filtering steps: (i) To enrich for potential promoter motifs, all candidate X-box sequence hits were required to locate within 1 kb upstream or downstream of position +1 of the start codon ATG of a given gene. (ii) We compared side-by-side the output of the search efforts employing canonical versus variant X-box query sequence PWMs. (iii) We compared the output lists of candidate X-box motifs and corresponding genes with *C. elegans* gene lists extracted from Wormbase (WS235; www.wormbase.org) where the required gene expression pattern is “ciliated neuron or labial neuron or inner labial neuron or IL2 neuron” (see also Wang *et al*., 2015). The resulting lists of high-confidence candidate X-box motifs and corresponding genes are presented in Tables EV4-EV6, all of which contain large numbers of novel candidate members of the DAF-19M regulatory subroutine in IL2 neurons.

## Acknowledgements

We thank Jiseon Lim and Eunkyeong Kim for helping with the bioinformatics-based sequence motif searches for the X-box motif variant. Worm expression vectors were kindly provided by Andrew Fire. Mutant worm strains were kindly provided by the *Caenorhabditis* Genetics Center (USA) and the National BioResource Project (Japan), Elizabeth De Stasio, H. Robert Horvitz, Darrell J. Killian, and Douglas S. Portman.

P. Swoboda acknowledges grant support from the following sources: Swedish Research Council Project grant, Swedish Research Council Equipment grant (Union Biometrica Worm Sorter), Swedish Research Council Sweden-Korea Exchange Program, STINT Organization Sweden-Korea Collaboration Program, Torsten Söderberg Foundation, Åhlén Foundation, OE & Edla Johansson Foundation, Karolinska Institute Strategic Neurosciences Program.

Research in the J. Lee laboratory was supported by the Korea-Sweden Research Cooperation through the National Research Foundation of Korea (NRF) (NRF-2016K1A3A1A47921615), the STINT Organization Korea-Sweden Collaboration Program, and the Basic Science Research Program through the National Research Foundation of Korea (NRF) (NRF-2019R1A6A1A10073437).

## Author contributions

Project conceptualization and planning: S. Ahn, P. Swoboda, J. Lee

Experimentation and methodology: S. Ahn, H. Yang, S. Son, D. Park, J. Lee

Data analysis: S. Ahn, H. Yang, S. Son, D. Park, P. Swoboda, J. Lee

Critical resources and reagents: S. Ahn, H. Yang, D. Park, J. Lee

Writing and illustrations – draft: S. Ahn, H. Yim, P. Swoboda, J. Lee

Writing and illustrations – editing and review: S. Ahn, H. Yim, P. Swoboda, J. Lee

Supervision and project management: P. Swoboda, J. Lee

Funding acquisition: P. Swoboda, J. Lee

## Expanded View Legends (Figures)

**Figure Expanded View 1** *daf-19m* regulates *klp-6* expression in IL2 neurons at post-embryonic stages, but is not important for the development of IL2 neurons.

**A** Confocal images of animals with an *Is* [*klp-6p::gfp*] integrated transgene in different *daf-19* mutant backgrounds: control –*jlIs1900*, the integrated transgene carrying *klp-6p::gfp* and other markers not relevant here; *daf-19(m86)* – the canonical null mutant; *daf-19(rh1024)* – a transposon *Tc1* derived mutant; *daf-19(tm5562)* – a *daf-19a/b* isoform specific mutant. Both *daf-19(m86)* and *daf-19(rh1024)*affect all *daf-19* isoforms including *daf-19m*, while in *daf-19(tm5562)* the *daf-19m* isoform remains intact. In every image, the dotted line outlines the shape of the worm. IL2 neurons are indicated (arrows). Scale bars are 20 μm.

**B** Confocal image of *Ex*[*F28A12.3p::gfp*] transgenic animals in a *daf-19(rh1024)* mutant background. *F28A12.3* is an IL2 neuron specific gene whose expression is largely *daf-19* independent (Phirke et al, 2011). The dotted line outlines the shape of the worm. IL2 neurons are indicated (arrows). Scale bars are 20 μm.

**C** Confocal images of *Ex[daf-19mp::gfp* and *klp-6p::mCherry]* transgenic animals in a wild-type N2 background at the L1 and dauer stages. Both stages are early juvenile developmental stages. In both images, the dotted line outlines the shape of the worm. IL2 neurons are indicated (arrows). Scale bars are 20 μm.

**Figure Expanded View 2** The X-box motif variant residing in the *klp-6* promoter controls *klp-6* expression not only in IL2 neurons but also in male specific neurons.

**A** In *C. elegans* the canonical X-box promoter motif, an imperfect inverted repeat, consists of a 6-nt 5’ half site (blue), a 2-nt spacer most often AT (black), and a 6-nt 3’ half site (orange). *klp-6* promoter sequence stretches strongly resembling these 5’ or 3’ half sites, respectively, are indicated. Distances are given as upstream of the *klp-6* start codon ATG.

**B** The X-box motif variant with a single nucleotide spacer (T) is conserved in the promoter of the IL2 neuron gene *klp-6* in *C. elegans* and in *klp-6* orthologs in other *Caenorhabditis* species: *Cbr* –*C. briggsae, Cbn* –*C. brenneri*. Blue – X-box 5’half site, black – single nucleotide spacer (T), orange – X-box 3’ half site. Distances are given as upstream of the *klp-6* start codon ATG.

**C** Confocal images of *Ex* [*klp-6p::gfp*] transgenic – male – animals carrying the indicated *klp-6* promoter truncation construct in a wild-type N2 background identified the relevant *klp-6* promoter region that controls both IL2 and male specific neuron expression. In every image, the dotted line outlines the shape of the worm. IL2 and CEM neurons are indicated in the male head (arrows). HOB and RnB neurons are indicated in the male tail (arrows). Scale bars are 20 μm.

**D** Confocal images of transgenic – male – animals carrying a tandem repeat of a *klp-6* X-box motif variant minimal promoter region fused to GFP. The tandem repeat used consists of an extended X-box motif variant (−628, −590). In both images, the dotted line outlines the shape of the worm. IL2 and CEM neurons are indicated in the male head (arrows). HOB and RnB neurons are indicated in the male tail (arrows). *klp-6p::mCherry* was used as the relevant neuron specific marker. Scale bars are 20 μm.

**Figure Expanded View 3** The X-box motif variants residing in the *cwp-4* promoter control *cwp-4* expression not only in IL2 neurons but also in male specific neurons.

**A** The X-box motif variant is conserved in the promoter of the IL2 neuron gene *cwp-4* in *C. elegans* and in *cwp-4* orthologs in other *Caenorhabditis* species: *Cbr* – *C. briggsae, Cre* – *C. remanei, Cbn* – *C. brenneri, Cjp* – *C. japonica*. Blue – X-box 5’ half site, black – spacer nucleotide, orange – X-box 3’ half site. Distances are given as upstream of the *cwp-4* start codon ATG.

**B** Confocal images of Ex[*cwp-4p::gfp*] transgenic – male – animals carrying the indicated *cwp-4* promoter mutation construct in a wild-type N2 background identified the relevant *cwp-4* promoter region that controls both IL2 and male specific neuron expression. The *cwp-4* promoter mutation consisted of substituting both X-box motif variant sequences (−90 to −78 and −68 to −56) with GGATCC C GGATCC. In every image, the dotted line outlines the shape of the worm. IL2 and CEM neurons are indicated in the male head (arrows). HOB and RnB neurons are indicated in the male tail (arrows). *klp-6p::mCherry* was used as the relevant neuron specific marker. Scale bars are 20 μm.

**C** Confocal images of Ex[*cwp-4p::gfp* and *klp-6p::mCherry*] transgenic animals in a wild-type N2 background at the L1 and dauer stages. Both stages are early juvenile developmental stages. In both images, the dotted line outlines the shape of the worm. IL2 neurons are indicated (arrows). Scale bars are 20 μm.

**Figure Expanded View 4** Schematics of **A** *daf-19* gene organization and constructs for *daf-19* isoform specific rescue experiments; and **B** *daf-19m* isoform promoter variants (wild type and specific substitutions) for investigating candidate IL2 neuron terminal selector protein binding sites.

**A** All the constructs for *daf-19* isoform specific rescue experiments are genomic DNA based with the exception of *pGG67. pGG67* is a fusion construct consisting of genomic DNA (exons 1-3) and complementary DNA (exons 3-12) for *daf-19a* isoform specific rescue (Senti & Swoboda, 2008). *pGG14* is a genomic DNA construct for *daf-19c* isoform specific rescue (Senti & Swoboda, 2008). *pJL1920* is a genomic DNA construct for *daf-19c* isoform specific rescue with the deletion of the HOB/RnB element (Wang et al, 2010). *pJL1921* is a genomic DNA construct for *daf-19m* isoform specific rescue that includes a 5’ fusion of the IL2/CEM enhancer element (Wang et al, 2010).

**B** The *daf-19m* promoter contains candidate binding sites for IL2 neuron terminal selector proteins, UNC-86 and CFI-1. Mutagenesis constructs eliminate candidate binding sites for UNC-86 (Substitution −761, –739), for both UNC-86 and CFI-1(Substitution −601, −566), and for CFI-1 (Substitution −132, −116).

## Expanded View Legends (Tables)

**Table Expanded View 1** A yeast-1-hybrid experiment with the isolated *klp-6 cis*-regulatory element as bait confirmed DAF-19 as its binding protein. We list the identity and numbers of yeast-1-hybrid clones that showed positive interaction between bait sequence and protein, as translated from a *C. elegans* cDNA library. All clones were examined and confirmed by PCR. In addition, some clones were sequenced for detailed binding site information.

**Table Expanded View 2** The list of *C. elegans* genes used for constructing a position weight matrix (PWM) for the canonical X-box motif. All genes were previously shown to harbor an X-box motif in their promoters (Blacque et al, 2005; Efimenko et al, 2005; Chen et al, 2006; Burghoorn et al, 2012). Available gene orthologs in other *Caenorhabditis* species (*C. briggsae* and *C. remanei*) were extracted from Wormbase (WS235; www.wormbase.org). The column Position describes the distance of the X-box motif upstream of position +1 of the start codon ATG.

**Table Expanded View 3** The list of *C. elegans* genes used for constructing a position weight matrix (PWM) for candidate X-box motif variants. Some of the genes were previously shown to harbor an X-box motif in their promoters (Blacque et al, 2005; Efimenko et al, 2005; Chen et al, 2006; Burghoorn et al, 2012), whereby we have added a number of genes expressed and functional in IL2 neurons including *klp-6, cwp-4, tat-6*, and *spg-20*. Available gene orthologs in other *Caenorhabditis* species (*C. briggsae*, *C. remanei*, *C. brenneri*) were extracted from Wormbase (WS235; www.wormbase.org). The column Position describes the distance of the X-box motif upstream of position +1 of the start codon ATG.

**Table Expanded View 4** A list of candidates for DAF-19M downstream target genes that harbor X-box motif variant hits with a single nucleotide spacer (candidate 13 nt X-box hits). Searches in *C. elegans* were carried out genome-wide. All sequence hits were extracted using FIMO (Find Individual Motif Occurrences; http://meme-suite.org/tools/fimo) based on a position weight matrix (Table EV3) that allowed for any single nucleotide spacer (N). The FIMO search parameter p-value was required to be smaller than 1E-04. All sequence hits listed were found to locate within 1 kb upstream or downstream of position +1 of the start codon ATG of the respective gene (column *Location)*. We have observed X-box motif sequence hits on both strands as indicated (*).

**Table Expanded View 5** A list of candidates for DAF-19M downstream target genes that harbor X-box motif variant hits with a double nucleotide spacer (candidate 14 nt X-box hits). Searches in *C. elegans* were carried out genome-wide. All sequence hits were extracted using FIMO (Find Individual Motif Occurrences; http://meme-suite.org/tools/fimo) based on a position weight matrix (Table EV3) that allowed for any double nucleotide spacer (NN). The FIMO search parameter p-value was required to be smaller than 1E-04. All sequence hits listed were found to locate within 1 kb upstream or downstream of position +1 of the start codon ATG of the respective gene (column *Location)*. We have observed X-box motif sequence hits on both strands as indicated (*). We have found X-box motif sequence hits located only on the (-) strand, while previous studies reported corresponding hits on the (+) strand, as indicated (**). In the case of the gene *nud-1*, Wormbase (WS235; www.wormbase.org) has revised the X-box promoter motif sequence from GTATCC AT GAAAAC (Efimenko et al, 2005) to GTATCC AT GGGAAC, as indicated (***). In this table X-box motif sequence hits that have been reported in previous studies are indicated in **bold**.

**Table Expanded View 6** A list of candidates for DAF-19M downstream target genes that harbor X-box motif variant hits with a triple nucleotide spacer (candidate 15 nt X-box hits). Searches in *C. elegans* were carried out genome-wide. All sequence hits were extracted using FIMO (Find Individual Motif Occurrences; http://meme-suite.org/tools/fimo) based on a position weight matrix (Table EV3) that allowed for any triple nucleotide spacer (NNN). The FIMO search parameter p-value was required to be smaller than 1E-04. All sequence hits listed were found to locate within 1 kb upstream or downstream of position +1 of the start codon ATG of the respective gene (column *Location)*. We have observed X-box motif sequence hits on both strands as indicated (*).

**Table Expanded View 7** List of *C. elegans* strains used in this study. The wild-type strain is Bristol N2. All strains used and listed in this Table are derived from this wild-type N2 background. The strains LJ896, LJ897 and LJ898 are marked with a single asterisk (*): they carry mutations that were originally received from other sources in a *him-5* mutant background (*cwp-4:* Douglas Portman lab; *daf-19:* Bob Horvitz lab; *klp-6:* CGC). We have removed the *him-5* mutant background by outcrossing with wild-type N2. The strain LJ899 is marked with a double asterisk (**): it carries the *cil-7(tm5848)*mutant allele after outcrossing with wild-type N2. We have originally received *cil-7(tm5848)* un-outcrossed from the National BioResource Project (Japan).

## References

1. Altun-Gultekin Z, Andachi Y, Tsalik EL, Pilgrim D, Kohara Y, Hobert O (2001) A regulatory cascade of three homeobox genes, ceh-10, ttx-3 and ceh-23, controls cell fate specification of a defined interneuron class in C. elegans. Development (Cambridge, England) 128: 1951–1969

2. Anvarian Z, Mykytyn K, Mukhopadhyay S, Pedersen LB, Christensen ST (2019) Cellular signalling by primary cilia in development, organ function and disease. Nat Rev Nephrol 15: 199–219

3. Barr MM, Garcia LR (2006) Male mating behavior. WormBook: the online review of C elegans biology: 1–11

4. Barr MM, Garcia LR, Portman DS (2018) Sexual Dimorphism and Sex Differences in Caenorhabditis elegans Neuronal Development and Behavior. Genetics 208: 909–935

5. Bhuckory S, Kays JC, Dennis AM (2019) In Vivo Biosensing Using Resonance Energy Transfer. Biosensors (Basel) 9

6. Blacque OE, Perens EA, Boroevich KA, Inglis PN, Li C, Warner A, Khattra J, Holt RA, Ou G, Mah AK et al (2005) Functional genomics of the cilium, a sensory organelle. Current biology: CB 15: 935–941

7. Brenner S (1974) The genetics of Caenorhabditis elegans. Genetics 77: 71–94

8. Burghoorn J, Piasecki BP, Crona F, Phirke P, Jeppsson KE, Swoboda P (2012) The in vivo dissection of direct RFX-target gene promoters in C. elegans reveals a novel cis-regulatory element, the C-box. Developmental biology 368: 415–426

9. Butcher RA, Fujita M, Schroeder FC, Clardy J (2007) Small-molecule pheromones that control dauer development in Caenorhabditis elegans. Nature chemical biology 3: 420–422

10. Cassada RC, Russell RL (1975) The dauerlarva, a post-embryonic developmental variant of the nematode Caenorhabditis elegans. Developmental biology 46: 326–342

11. Chen N, Mah A, Blacque OE, Chu J, Phgora K, Bakhoum MW, Newbury CR, Khattra J, Chan S, Go A et al (2006) Identification of ciliary and ciliopathy genes in Caenorhabditis elegans through comparative genomics. Genome biology 7: R126

12. Choksi SP, Lauter G, Swoboda P, Roy S (2014) Switching on cilia: transcriptional networks regulating ciliogenesis. Development (Cambridge, England) 141: 1427–1441

13. Colbert HA, Smith TL, Bargmann CI (1997) OSM-9, a novel protein with structural similarity to channels, is required for olfaction, mechanosensation, and olfactory adaptation in Caenorhabditis elegans. The Journal of neuroscience: the official journal of the Society for Neuroscience 17: 8259–8269

14. De Stasio EA, Mueller KP, Bauer RJ, Hurlburt AJ, Bice SA, Scholtz SL, Phirke P, Sugiaman-Trapman D, Stinson LA, Olson HB et al (2018) An Expanded Role for the RFX Transcription Factor DAF-19, with Dual Functions in Ciliated and Non-ciliated Neurons. Genetics

15. Efimenko E, Bubb K, Mak HY, Holzman T, Leroux MR, Ruvkun G, Thomas JH, Swoboda P (2005) Analysis of xbx genes in C. elegans. Development (Cambridge, England) 132: 1923–1934

16. Emery P, Durand B, Mach B, Reith W (1996a) RFX proteins, a novel family of DNA binding proteins conserved in the eukaryotic kingdom. Nucleic acids research 24: 803–807

17. Emery P, Strubin M, Hofmann K, Bucher P, Mach B, Reith W (1996b) A consensus motif in the RFX DNA binding domain and binding domain mutants with altered specificity. Molecular and cellular biology 16: 4486–4494

18. Etchberger JF, Flowers EB, Poole RJ, Bashllari E, Hobert O (2009) Cis-regulatory mechanisms of left/right asymmetric neuron-subtype specification in C. elegans. Development (Cambridge, England) 136: 147–160

19. Etchberger JF, Lorch A, Sleumer MC, Zapf R, Jones SJ, Marra MA, Holt RA, Moerman DG, Hobert O (2007) The molecular signature and cis-regulatory architecture of a C. elegans gustatory neuron. Genes & development 21: 1653–1674

20. Gajiwala KS, Chen H, Cornille F, Roques BP, Reith W, Mach B, Burley SK (2000) Structure of the winged-helix protein hRFX1 reveals a new mode of DNA binding. Nature 403: 916–921

21. Gordon PM, Hobert O (2015) A competition mechanism for a homeotic neuron identity transformation in C. elegans. Developmental cell 34: 206–219

22. Hobert O (2002) PCR fusion-based approach to create reporter gene constructs for expression analysis in transgenic C. elegans. Biotechniques 32: 728–730

23. Hobert O (2008) Regulatory logic of neuronal diversity: terminal selector genes and selector motifs. Proceedings of the National Academy of Sciences of the United States of America 105: 20067–20071

24. Hobert O (2011) Regulation of terminal differentiation programs in the nervous system. Annual review of cell and developmental biology 27: 681–696

25. Hobert O (2016) Terminal Selectors of Neuronal Identity. Current topics in developmental biology 116: 455–475

26. Hurd DD, Miller RM, Nunez L, Portman DS (2010) Specific alpha-and beta-tubulin isotypes optimize the functions of sensory Cilia in Caenorhabditis elegans. Genetics 185: 883–896

27. Inglis PN, Ou G, Leroux MR, Scholey JM (2007) The sensory cilia of Caenorhabditis elegans. WormBook: the online review of C elegans biology: 1–22

28. Jeong PY, Jung M, Yim YH, Kim H, Park M, Hong E, Lee W, Kim YH, Kim K, Paik YK (2005) Chemical structure and biological activity of the Caenorhabditis elegans dauer-inducing pheromone. Nature 433: 541–545

29. Jolma A, Yan J, Whitington T, Toivonen J, Nitta KR, Rastas P, Morgunova E, Enge M, Taipale M, Wei G et al (2013) DNA-binding specificities of human transcription factors. Cell 152: 327–339

30. Lai HT, Chiang CM (2013) Bimolecular Fluorescence Complementation (BiFC) Assay for Direct Visualization of Protein-Protein Interaction in vivo. Bio Protoc 3

31. Laurençon A, Dubruille R, Efimenko E, Grenier G, Bissett R, Cortier E, Rolland V, Swoboda P, Durand B (2007) Identification of novel regulatory factor X (RFX) target genes by comparative genomics in Drosophila species. Genome biology 8: R195

32. Lee D, Lee H, Choi M-k, Park S, Lee J (2015) Nictation Assays for Caenorhabditis and Other Nematodes. Bio-protocol 5: e1433

33. Lee D, Yang H, Kim J, Brady S, Zdraljevic S, Zamanian M, Kim H, Paik YK, Kruglyak L, Andersen EC et al (2017) The genetic basis of natural variation in a phoretic behavior. Nature communications 8: 273

34. Lee H, Choi MK, Lee D, Kim HS, Hwang H, Kim H, Park S, Paik YK, Lee J (2012) Nictation, a dispersal behavior of the nematode Caenorhabditis elegans, is regulated by IL2 neurons. Nat Neurosci 15: 107–112

35. Lemeille S, Paschaki M, Baas D, Morlé L, Duteyrat JL, Ait-Lounis A, Barras E, Soulavie F, Jerber J, Thomas J et al (2020) Interplay of RFX transcription factors 1, 2 and 3 in motile ciliogenesis. Nucleic acids research

36. Maguire JE, Silva M, Nguyen KC, Hellen E, Kern AD, Hall DH, Barr MM (2015) Myristoylated CIL-7 regulates ciliary extracellular vesicle biogenesis. Molecular biology of the cell 26: 2823–2832

37. Mello CC, Kramer JM, Stinchcomb D, Ambros V (1991) Efficient gene transfer in C.elegans: extrachromosomal maintenance and integration of transforming sequences. The EMBO journal 10: 3959–3970

38. Miller RM, Portman DS (2010) A latent capacity of the C. elegans polycystins to disrupt sensory transduction is repressed by the single-pass ciliary membrane protein CWP-5. Disease models & mechanisms 3: 441–450

39. Mitchison HM, Valente EM (2017) Motile and non-motile cilia in human pathology: from function to phenotypes. J Pathol 241: 294–309

40. Morsci NS, Barr MM (2011) Kinesin-3 KLP-6 regulates intraflagellar transport in male-specific cilia of Caenorhabditis elegans. Current biology: CB 21: 1239–1244

41. Mukhopadhyay S, Lu Y, Qin H, Lanjuin A, Shaham S, Sengupta P (2007) Distinct IFT mechanisms contribute to the generation of ciliary structural diversity in C. elegans. The EMBO journal 26: 2966–2980

42. Nachury MV, Mick DU (2019) Establishing and regulating the composition of cilia for signal transduction. Nature reviews Molecular cell biology 20: 389–405

43. Ouellet J, Li S, Roy R (2008) Notch signalling is required for both dauer maintenance and recovery in C. elegans. Development (Cambridge, England) 135: 2583–2592

44. Peden EM, Barr MM (2005) The KLP-6 kinesin is required for male mating behaviors and polycystin localization in Caenorhabditis elegans. Current biology: CB 15: 394–404

45. Perkins LA, Hedgecock EM, Thomson JN, Culotti JG (1986) Mutant sensory cilia in the nematode Caenorhabditis elegans. Developmental biology 117: 456–487

46. Phirke P, Efimenko E, Mohan S, Burghoorn J, Crona F, Bakhoum MW, Trieb M, Schuske K, Jorgensen EM, Piasecki BP et al (2011) Transcriptional profiling of C. elegans DAF-19 uncovers a ciliary base-associated protein and a CDK/CCRK/LF2p-related kinase required for intraflagellar transport. Developmental biology 357: 235–247

47. Piasecki BP, Burghoorn J, Swoboda P (2010) Regulatory Factor X (RFX)-mediated transcriptional rewiring of ciliary genes in animals. Proceedings of the National Academy of Sciences of the United States of America 107: 12969–12974

48. Portman DS, Emmons SW (2004) Identification of C. elegans sensory ray genes using whole-genome expression profiling. Developmental biology 270: 499–512

49. Rand JB (1989) Genetic analysis of the cha-1-unc-17 gene complex in Caenorhabditis. Genetics 122: 73–80

50. Reed EM, Wallace HR (1965) Leaping Locomotion by an Insect-parasitic Nematode. Nature 206: 210–211

51. Reiter JF, Leroux MR (2017) Genes and molecular pathways underpinning ciliopathies. Nature reviews Molecular cell biology 18: 533–547

52. Schroeder NE, Androwski RJ, Rashid A, Lee H, Lee J, Barr MM (2013) Dauer-specific dendrite arborization in C. elegans is regulated by KPC-1/Furin. Current biology: CB 23: 1527–1535

53. Senti G, Swoboda P (2008) Distinct isoforms of the RFX transcription factor DAF-19 regulate ciliogenesis and maintenance of synaptic activity. Molecular biology of the cell 19: 5517–5528

54. Shaham S (2010) Chemosensory organs as models of neuronal synapses. Nat Rev Neurosci 11: 212–217

55. Starich TA, Herman RK, Kari CK, Yeh WH, Schackwitz WS, Schuyler MW, Collet J, Thomas JH, Riddle DL (1995) Mutations affecting the chemosensory neurons of Caenorhabditis elegans. Genetics 139: 171–188

56. Sugiaman-Trapman D, Vitezic M, Jouhilahti EM, Mathelier A, Lauter G, Misra S, Daub CO, Kere J, Swoboda P (2018) Characterization of the human RFX transcription factor family by regulatory and target gene analysis. BMC genomics 19: 181

57. Sulston JE, Schierenberg E, White JG, Thomson JN (1983) The embryonic cell lineage of the nematode Caenorhabditis elegans. Developmental biology 100: 64–119

58. Swoboda P, Adler HT, Thomas JH (2000) The RFX-type transcription factor DAF-19 regulates sensory neuron cilium formation in C. elegans. Mol Cell 5: 411–421

59. Wang J, Kaletsky R, Silva M, Williams A, Haas LA, Androwski RJ, Landis JN, Patrick C, Rashid A, Santiago-Martinez D et al (2015) Cell-Specific Transcriptional Profiling of Ciliated Sensory Neurons Reveals Regulators of Behavior and Extracellular Vesicle Biogenesis. Current biology: CB 25: 3232–3238

60. Wang J, Schwartz HT, Barr MM (2010) Functional specialization of sensory cilia by an RFX transcription factor isoform. Genetics 186: 1295–1307

61. Ward S, Thomson N, White JG, Brenner S (1975) Electron microscopical reconstruction of the anterior sensory anatomy of the nematode Caenorhabditis elegans. J Comp Neurol 160: 313–337

62. Wells KL, Rowneki M, Killian DJ (2015) A splice acceptor mutation in C. elegans daf-19/Rfx disrupts functional specialization of male-specific ciliated neurons but does not affect ciliogenesis. Gene 559: 196–202

63. White JG, Southgate E, Thomson JN, Brenner S (1986) The structure of the nervous system of the nematode Caenorhabditis elegans. Philos Trans R Soc Lond B Biol Sci 314: 1–340

64. Xie Y, Moussaif M, Choi S, Xu L, Sze JY (2013) RFX transcription factor DAF-19 regulates 5-HT and innate immune responses to pathogenic bacteria in Caenorhabditis elegans. PLoS genetics 9: e1003324

65. Zhang F, Bhattacharya A, Nelson JC, Abe N, Gordon P, Lloret-Fernandez C, Maicas M, Flames N, Mann RS, Colon-Ramos DA et al (2014) The LIM and POU homeobox genes ttx-3 and unc-86 act as terminal selectors in distinct cholinergic and serotonergic neuron types. Development (Cambridge, England) 141: 422–435

66. Zhang H, Yue X, Cheng H, Zhang X, Cai Y, Zou W, Huang G, Cheng L, Ye F, Kang L (2018) OSM-9 and an amiloride-sensitive channel, but not PKD-2, are involved in mechanosensation in C. elegans male ray neurons. Scientific reports 8: 7192

